# ecDNA replication is disorganised and vulnerable to replication stress

**DOI:** 10.1101/2025.03.22.644567

**Authors:** Jedrzej J. Jaworski, Pauline L. Pfuderer, Pawel Czyz, Gianluca Petris, Michael A. Boemo, Julian E. Sale

## Abstract

Extrachromosomal DNA (ecDNA) is a critical driver of cancer progression, contributing to tumour growth, evolution, and therapeutic resistance through oncogene amplification. Despite its significance, the replication of ecDNA remains poorly understood. In this study, we investigated the replication dynamics of ecDNA using high-resolution replication timing analysis (Repli-seq) and DNAscent, a method for measuring origin firing and replication fork movement based on ultra-long read Oxford Nanopore Sequencing, that we applied to both bulk DNA and to ecDNA isolated with FACS-based Isolation of Native ecDNA (FINE), a new method for isolating intact, chromatinised ecDNA without DNA or protein digestion. We demonstrate that ecDNA in the COLO 320DM colorectal cancer cell line exhibits largely asynchronous replication throughout the S phase, contrasting with the conserved replication timing of the corresponding normal linear chromosomal DNA in RPE-1 cells and the chromosomally reintegrated ecDNA in COLO 320HSR, which forms a homogeneously staining region. Replication origins on ecDNA are redistributed, and replication forks exhibit reduced velocity and increased stalling, particularly near the c-*MYC* oncogene. Under replication stress induced by hydroxyurea treatment, ecDNA replication is further compromised, leading to altered origin activation, reduced fork velocity and eventual ecDNA depletion from cells. Our findings reveal fundamental differences in the replication dynamics of ecDNA, providing insights that could inform the development of therapies targeting ecDNA-associated oncogene amplification in cancer.

## Introduction

Oncogene amplification is a critical driver of cancer progression, contributing to tumour growth, evolution and resistance to therapy. Traditionally, oncogene amplification has been associated with copy number increase on linear chromosomes, observed as homogeneously staining regions (HSRs) in chromosome G-banding (1). However, recent studies have highlighted an alternative and highly potent mechanism of oncogene amplification through extrachromosomal DNA (ecDNA) (2, 3).

ecDNA molecules comprise large, double stranded DNA fragments averaging between 1-3 Mb in size (2) that have been detected in approximately 17 % of all cancers (4). These circular DNA elements frequently harbour oncogenes or immunomodulatory genes (3, 5, 6), driving the aggressive behaviour of tumours. ecDNA-based oncogenes are highly transcribed, even when adjusted for their high copy number, which often exceeds 100 copies per cell (7).

Unlike canonical chromosomes, ecDNA lacks centromeres, leading to its uneven segregation during cell division (8, 9). This unique flexibility of ecDNA contributes to the rapid evolution of tumours by promoting intratumour genetic heterogeneity (9) allowing cancer cells to rapidly adapt to selective pressures, including therapeutic interventions (9–13). Consequently, the presence of ecDNA is associated with poorer outcomes, including reduced survival rates compared to tumours without such amplifications (4, 5).

Despite the recognised importance of ecDNA in cancer, much remains unknown about its DNA replication dynamics. While ecDNA clearly replicates (14, 15), and likely only once per cell cycle during S phase (16), details such as the precise timing of replication, the distribution of replication origins, replication fork velocity and the frequency of replication stalling remain poorly understood. For instance, it is unclear whether the replication origins and replication timing found in a chromosomal segment are preserved when present in ecDNA. Moreover, ecDNA is characterised by elevated replication stress (17), and its loss appears to be linked primarily to a further increase in this stress followed by micronuclei-mediated ecDNA elimination, particularly following treatment with hydroxyurea (HU) (18–21). A more complete understanding of ecDNA replication may enable the development of strategies to control ecDNA copy number and, consequently, oncogenic drive in certain cancers.

To address these questions, we combine assessment of the replication timing programme using Repli-seq (22, 23) with DNAscent (24), a method for determining the velocity and direction of the nascent DNA synthesis in single DNA molecules, applying Oxford Nanopore Technologies (ONT) long-read sequencing of COLO 320DM cells and matched ecDNA-negative controls. Further, to directly detect replicating ecDNA, we introduce FINE (FACS-based Isolation of Native ecDNA), a method for isolating largely intact, chromatinised ecDNA without requiring DNA or protein digestion. We show that replication timing programme in ecDNA is significantly disrupted. Further, replication in ecDNA appears more sensitive to stress induced by depletion of ribonucleotide pools with HU than chromosomal DNA, which leads to replication stress-induced ecDNA loss.

## Materials & Methods

### Cell Culture

Human colorectal adenocarcinoma cell lines COLO 320DM and COLO 320HSR were obtained from the American Type Culture Collection (ATCC) and maintained in RPMI-1640 medium (Gibco) supplemented with 10% (vol/vol) fetal bovine serum (FBS; Sigma-Aldrich). hTERT-immortalized retinal pigment epithelial cells (RPE-1) were also sourced from ATCC and cultured in a 1:1 mixture of Dulbecco’s Modified Eagle’s Medium and Ham’s F12 (DMEM, Gibco) supplemented with 10% (vol/vol) FBS. All cell lines were incubated in a humidified atmosphere of 5% CO2 at 37°C. Cell lines were regularly tested for mycoplasma contamination.

### Metaphase Chromosome Spreads

Cells were arrested in metaphase by treatment with either 0.1 µg/ml KaryoMAX Colcemid solution in PBS (Gibco) for RPE-1 cells, or 0.4 µM nocodazole (Sigma-Aldrich) for COLO 320DM and COLO 320HSR cells, for 3 h at 37°C. After arrest, cells were washed with PBS and gently resuspended in a pre-warmed 75 mM potassium chloride (KCl) hypotonic solution, followed by a 20 min incubation at 37°C to swell the cells.

Cells were pre-fixed by adding 10% volume of freshly prepared Carnoy’s fixative (3:1 methanol:acetic acid, vol/vol), then washed twice with ice-cold Carnoy’s fixative and stored overnight at −20°C. The following day, the cell suspension was dropped from a height of approximately 50 cm onto ice-cold glass slides held at a 45° angle. Slides were matured by incubation at 65°C for 1 h, followed by washing in 2× saline-sodium citrate (SSC) buffer. Chromosomes were stained with 1 µg/ml Hoechst 33258 (Sigma-Aldrich) for visualisation.

### Fluorescence In Situ Hybridization (FISH)

Fixed metaphase spreads were washed in 2× saline-sodium citrate (SSC) buffer and sequentially dehydrated ethanol series (70%, 85%, 100%) for 2 min each. Fluorescence in situ hybridization (FISH) probes (Empire Genomics; green CHR08-10-GR for Chromosome 8 and orange MYC-20-OR for c-Myc) were applied to the slides, which were then sealed with rubber cement. Denaturation was performed on a hot plate at 75°C for 7 min, followed by hybridization for 16 h in a humidified chamber at 37°C.

After hybridization, coverslips were removed, and slides were washed in 0.3% Igepal CA-630 (Sigma-Aldrich) in 0.4× SSC at 73°C for 2 min, followed by a wash in 0.1% Igepal CA-630 in 2× SSC at room temperature for 2 min. The slides were stained with 1 µg/ml Hoechst 33258 (Sigma-Aldrich) and washed again in 2× SSC before being mounted with ProLong Diamond Antifade Mountant (Thermo Fisher Scientific) and covered with a coverslip.

FISH images were captured using a Zeiss LSM780 confocal microscope with a 63× objective. Image analysis was performed using Fiji (ImageJ-based software) (25).

### Chromosome Flow Cytometry and Sorting

Actively dividing COLO 320DM and COLO 320HSR cells were synchronised in mitosis by treating with 0.4 µM nocodazole (Sigma-Aldrich) for 13 h. For RPE-1 cells, 100 µl of Colcemid (Gibco KaryoMAX Colcemid Solution in PBS) was added to 30 ml of culture medium. Following mitotic arrest, cells were harvested by mitotic shake-off and collected by centrifugation at 400g for 5 min at room temperature. Cells were resuspended in the medium containing nocodazole and incubated with 0.1 µg/ml Latrunculin B (Cambridge Bioscience CAY10010631) at 37°C for 1 h.

Cells were pelleted by centrifugation (400g, 5 min) and resuspended in a hypotonic solution containing 75 mM KCl, 10 mM MgSO_4_, 0.2 mM spermine, and 0.5 mM spermidine, pH 8.0. The cell suspension was incubated at room temperature for 20 min, followed by centrifugation at 300g for 5 min. Cells were then resuspended in ice-cold PAB buffer (150 mM Tris-HCl, pH 8.0, 0.25% Triton X-100, 20 mM KCl, 0.5 mM spermine, 1 mM EDTA, 5 mM EGTA). Chromosomes were released by gently flicking the tube, and the quality of chromosome release was checked by mixing 10 µl of sample with 1 µl of propidium iodide (PI) and visualising under a microscope.

The isolated chromosomes were stained with a combination of 5 µg/ml DAPI, 50 µg/ml Chromomycin A3, and 10 mM MgSO4. The staining reaction was carried out in a low volume (300 µl) for 1 h at room temperature, followed by dilution with 2–3 ml of PAB buffer. Samples were then incubated for an additional 30 min.

Chromosomes were analysed and sorted using a BD FACSAria Fusion Flow Cytometer with a 70-µm or 100-µm nozzle, set at a flow rate of < 25,000 events/sec. The 585 nm filter was used to detect Chromomycin A3 and 450 nm filter was used to detect DAPI. Sorted chromosomes were maintained in PBS containing 1 mM EDTA for further analysis.

### Whole-Genome Sequencing (WGS) and ecDNA Assembly

Genomic DNA was extracted from cultured cells using the Monarch Genomic DNA Purification Kit (New England Biolabs) according to the manufacturer’s protocol. DNA libraries were prepared using the NEBNext® Ultra™ II FS DNA Library Prep Kit for Illumina (New England Biolabs), following the manufacturer’s instructions. Library quantification and dilution were performed using the Qubit dsDNA High Sensitivity Quantification Assay (Thermo Fisher Scientific). Sequencing was carried out on an Illumina NextSeq 2000 platform.

The resulting FASTQ files were aligned to the human reference genome (hg38) using BWA-MEM. AmpliconArchitect (AA; https://github.com/virajbdeshpande/AmpliconArchitect; (26)) was employed to detect genomic amplifications. The output from AA was integrated with optical genome mapping data to facilitate the assembly of extrachromosomal DNA (ecDNA) structures using the Amplicon Classifier tool. Optical genome mapping was performed by the Wellcome Sanger Institute using the Saphyr® instrument (Bionano Genomics). Cell pellets were prepared following standard Bionano protocols. DNA was labelled by DLE-1 and de novo assembled using the Bionano Access software.

### Repli-seq analysis of DNA replication timing

Repli-seq was performed as previously described with modifications (23, 27). Briefly, actively proliferating cells were cultured in T175 flasks under standard conditions and pulse-labelled with 100 µM BrdU (Sigma-Aldrich) for 30 min at 37°C. After labelling, cells were washed twice with ice-cold PBS and fixed by adding 75% (vol/vol) ice-cold ethanol dropwise while vortexing gently. Cells were stored at −20°C for at least 16 h.

For FACS, fixed cells were washed with 1% (vol/vol) FBS in PBS and stained with a solution of propidium iodide (50 μg/ml, Sigma-Aldrich) and RNase A (20 μg/ml, Sigma-Aldrich) in PBS/1% FBS. After 30 min of incubation at room temperature in the dark, cells were filtered through a 37-μm nylon mesh and sorted by flow cytometry (BD FACSAria II) into five fractions representing different stages of S phase based on DNA content.

Genomic DNA was extracted from sorted cells using the Zymo Quick-DNA Microprep Kit (Zymo Research) following the manufacturer’s instructions. DNA was fragmented to an average size of 200 bp using a Covaris M220 ultrasonicator set to 75 W peak incident power, 10% duty cycle, 200 cycles per burst, for 260 seconds. Fragment size was verified using an Agilent Bioanalyzer 2100 with a high-sensitivity DNA chip.

Next, libraries were prepared using the NEBNext Ultra DNA Library Prep Kit for Illumina (NEB), according to the manufacturer’s instructions. BrdU-labeled DNA was immunoprecipitated by first denaturing the DNA at 95°C for 5 min, followed by incubation with an anti-BrdU antibody (BD Biosciences). Immunoprecipitated complexes were recovered by incubation with rabbit anti-mouse IgG and centrifugation. After washing, the samples were digested overnight with proteinase K (0.25 mg/ml) at 56°C. DNA was purified using the DNA Clean & Concentrator-5 kit (Zymo Research). The purified DNA was amplified, indexed, and prepared for sequencing using NEBNext Ultra II DNA Library Prep Kit for Illumina. Sequencing was performed on an Illumina NextSeq 2000 platform.

Sequencing reads were aligned to the human reference genome (GRCh38) using BWA-MEM. The aligned data were indexed, and PCR duplicates and reads with multiple alignments were removed using SAMtools (28). The resulting BAM files were normalized to reads per kilobase per million (RPKM) using bamCoverage. The normalized data were visualised in the Integrative Genomics Viewer (IGV) (29).

The statistical analysis of the synchronicity of the replication was achieved using Rao’s quadratic entropy estimate. We consider bins of length 10kB, to which reads are mapped and normalised between five fractions encompassing the S phase, giving a categorical distribution over five classes. As a measure of disorder, we use Rao’s quadratic entropy (30, 31) with a circular metric, which is more suitable for modelling cycling cells than alternatives not employing the temporal order of the categories, such as Shannon’s entropy (32). The amplified fragment of length 1.6 Mb hence contains 160 bins, each associated with a single Rao’s quadratic entropy estimate. We average this value over all bins, to associate a single estimate σ to each 1.6 Mb genomic fragment.

To assess the degree of extremity of the observed value, we calculate the empirical distribution of Rao’s quadratic entropy estimate over fragments of matched length chosen uniformly along the whole genome, with the exception of fragments of length 5 Mb from each end to avoid sequencing artifacts. To avoid correlations between closely-located fragments we ensure 10Mb spaces between subsequent fragments. This yields N = 273 fragments distributed along 22 chromosomes. After removing fragments containing bins without reads, we use N = 223 fragments of 1.6 Mb to estimate the empirical distribution. We compared the empirical distributions of RQE between three cell lines using two-sample Kolmogorov-Smirnov test. All three comparisons returned highly significant p-values (< 10^-19^ for HSR – DM; < 10^-8^ for RPE-1 – HSR; 0.003 for RPE-1 – DM), suggesting minor experiment-specific differences. However, qualitatively the distributions look similar (Figure 2B). To assess the extremity of the RQE value associated with the *MYC* fragment, we calculated the tail probabilities p = P(X >= X_c-*MYC*_). We obtain p_DM < 0.01 and p_HSR < 0.01, with p_RPE-1 = 0.23. Our results therefore suggest that the replication of the amplified fragment in COLO 320DM and COLO 320HSR proceeds in a less ordered fashion than for the large majority of the matching fragments in these cell lines.

### RT–qPCR

Quantitative PCR (qPCR) was employed to estimate copy number variations of ecDNA by targeting the c-*MYC* gene (TaqMan™ Copy Number Assay, Assay ID: Hs00292858_cn). The human RNase P gene (TaqMan™ Copy Number Reference Assay, RNase P; Applied Biosystems, Cat. No. 4403328) was used as the reference.

Reactions were prepared using TaqPath™ ProAmp™ Master Mix (Thermo Fisher Scientific, Cat. No. A30865) according to the manufacturer’s instructions. Each 20 µL reaction consisted of 10 µL of 2× TaqPath™ ProAmp™ Master Mix, 1 µL of 20× TaqMan™ Copy Number Assay, 1 µL of 20× TaqMan™ Copy Number Reference Assay (RNase P), and 2 µL of genomic DNA (1.6 ng). Amplification was conducted on an Viia7 (ThermoFisher) cycler under the following thermal cycling conditions: initial denaturation at 95°C for 10 minutes, followed by 40 cycles of 95°C for 15 seconds and 60°C for 60 seconds.

Relative copy number was calculated using the comparative Ct (ΔΔCt) method, normalising target Ct values to the reference RNase P gene and comparing them to a TK-6 sample with a known copy number. All reactions were performed in technical quadruplicate across three biological replicates.

### Replication Dynamics and Origin Analysis with DNAscent

Replication velocity and origins were analysed using DNAscent. Actively dividing cells in COLO 320DM-HU sample were treated with 50 µM hydroxyurea for 24 h to induce replication stress. After the optional treatment, cells were pulse-labelled with 50 µM EdU for 6 min, washed twice with warm PBS, and then treated with 50 µM BrdU for 6 min. Cells were washed again with warm PBS and incubated with 100 µM thymidine for 1 h. After a final PBS wash, fresh complete medium was added.

For PromethION (ONT) sequencing, cells were harvested 1 h after thymidine treatment, washed with ice-cold PBS containing 1% BSA, and flash-frozen in liquid nitrogen. Approximately 6 million cells were used for each experiment. DNA libraries were prepared for ultra-long sequencing using the Ultra-Long DNA Sequencing Kit (SQK-ULK1, ONT) and sequenced on a Nanopore PromethION platform. For one of the untreated DM samples (DM_untreated_rep2) adaptive sampling targeting chromosome 8 was performed (Supplementary Table 1). An initial attempt at enriching circles with pulse field gel electrophoresis resulted in a low N_50_ and no enrichment (Supplementary Table 1). N50 is calculated for reads passing the DNAscent quality criteria (minimum alignment length of 20 kb and minimum mapping quality of 20). Then, all reads are sorted by length in descending order, and the cumulative sum over all read lengths is calculated. To determine N50, the read where the cumulative sum up to that read is equal or greater than half of the overall cumulative sum, is determined, and this read’s length is the N50. An overview of all datasets is provided in Supplementary Table 1.

Alternatively, 30 min after cell labelling with BrdU and EdU, cells were arrested in mitosis by overnight treatment with 0.4 µM nocodazole. Following ecDNA release and FACS sorting (as described above), ecDNA was concentrated by centrifugation at 24,000g for 1 h at 4°C. DNA libraries were prepared using the Ligation Sequencing Kit (SQK-LSK110) and sequenced on a ONT MinION platform, following the manufacturer’s instructions.

Base calling of the raw sequencing files from ONT PromethION or MinION platform (R9.4.1 Nanopore, fast5 files available on ENA under accession PRJEB83636) was performed using GUPPY (version 5.0.16) with the configuration for R9-sequenced DNA (dna_r9.4.1_450bps_fast.cfg) and the output reads (fastq format) that passed and failed the base calling quality metrics were used for subsequent processing. Alignment of the reads to the hg38 reference genome (release GRCh38.p14) or our custom ecDNA reference map (see above) was performed with minimap2 (version 2.24). The output from minimap2 (sam format) was then converted to bam format and sorted and indexed using samtools (version 1.14). Next, DNAscent (version 3.1.2) subprogrammes index, detect, forkSense and bedgraph (visualisation) were run and replication forks and origins were visualised using a genome browser (Integrative Genomics Viewer, version 2.15.1).

### Calculation of replication fork speed and outlier removal

The DNAscent forkSense output (bed file format, available on ENA under accession PRJEB83636) was used for custom downstream processing using Python (version 3.8.2), starting with calculation of fork speed. Replication fork speed and stall scores from DNAscent were obtained as previously described (33). We only include replication forks where both the fork speed and stall score could be calculated for downstream analyses i.e., fork tracks were not near the end of the read. Reads that mapped to chromosome Y were removed as the cell line originates from the tumour of a female patient (34). For all fork speed figures in the main body, we apply outlier filtering based on the interquartile range (IQR). Outliers were removed based on the IQR. The 1st (Q1, 25%) and 3rd quartiles (Q3, 75%) were calculated, and the IQR was defined as the difference between Q3 and Q1. Outliers were identified as data points lying below the lower bound (Q1 - 1.5 * IQR) or above the upper bound (Q3 + 1.5 * IQR) and were excluded from further analysis. For the supplementary histograms of fork speed, no IQR filtering was applied to present the raw data (Supplementary Figure 5). A replicate comparison for the PromethION runs is provided in Supplementary Figure 6. Data cleaning, outlier removal, statistics and visualisations were performed using custom Python (version 3.11.5) scripts, except for circular plots which were created in R (see below).

### FACS enrichment analysis

To assess the enrichment factor achieved through FACS sorting, we compared the ratio of reads mapped to ecDNA and to the human reference genome, focusing only on reads that passed the DNAscent minimum quality criteria (minimum alignment length of 20 kb and minimum mapping quality of 20).

### Visualisation of ecDNA maps with origin densities, fork speeds and stall scores

Circular ecDNA visualisations were generated using the *circlize* package (version 0.4.16) (35) in R (version 4.3.3) using the IQR filtered replication forks as input for fork speed and stall scores and DNAscent forkSense origins for origin of replication analyses. In brief, a custom backbone was created using the AmpliconArchitect (26) generated ecDNA reference map. For origin of replication visualisations, origins from all biological replicates of COLO 320DM (‘DM’, 4 replicates), COLO 320HSR (‘HSR’, 4 replicates) or DM HU treated (‘DM-HU’, 3 replicates) were pooled. Origins were grouped into 50 kb segments and normalised by the number of reads in each segment to account for varying read depth within and across samples. The number of reads in each segment was obtained from the aligned bam file and only reads that passed the DNAscent quality criteria (minimum alignment length of 20 kb and minimum mapping quality of 20) were included for the read depth assessment. Aligned read length and mapping quality were extracted from the bam file using the *pysam* package (https://github.com/pysam-developers/pysam) in Python. For replication fork speed and stall scores, replication forks for each group (DM, HSR, DM-HU) from all replicates were pooled and grouped into 20 kb segments. Fork speed and stall score averages across all forks in each segment were calculated for visualisation. G-quadruplex scores for the ecDNA reference map were obtained using G4Hunter (36) with a window size of 25 and a threshold of 1.5.

## Results

### Molecular Characterisation and Isolation of ecDNA

To investigate ecDNA replication, we used the colorectal cancer cell line COLO 320DM (34), which carries an amplification of the c-*MYC* locus on multiple ecDNAs. A cell line (COLO 320HSR) derived from the same patient tumour provides an interesting comparison in that it harbours linear amplifications of a region of chromosome 8 containing c-*MYC* on another chromosome (34). We also used an unrelated, immortalised, but untransformed cell line RPE-1 as a control (37) (Figure 1A). We characterised the ecDNA structure in our isolate of COLO 320DM, using AmpliconArchitect (26) and AmpliconReconstructor (38) algorithms that integrate optical genome mapping with whole-genome short-read sequencing (Figure 1B). COLO 320DM contained a heterogeneous population of ecDNA containing the fragment of chromosome 8 surrounding c-*MYC* oncogene, and for our analysis we selected the predominant 1.5Mb species obtained using AmpliconArchitect (Figure 1B).

**Figure 1.**
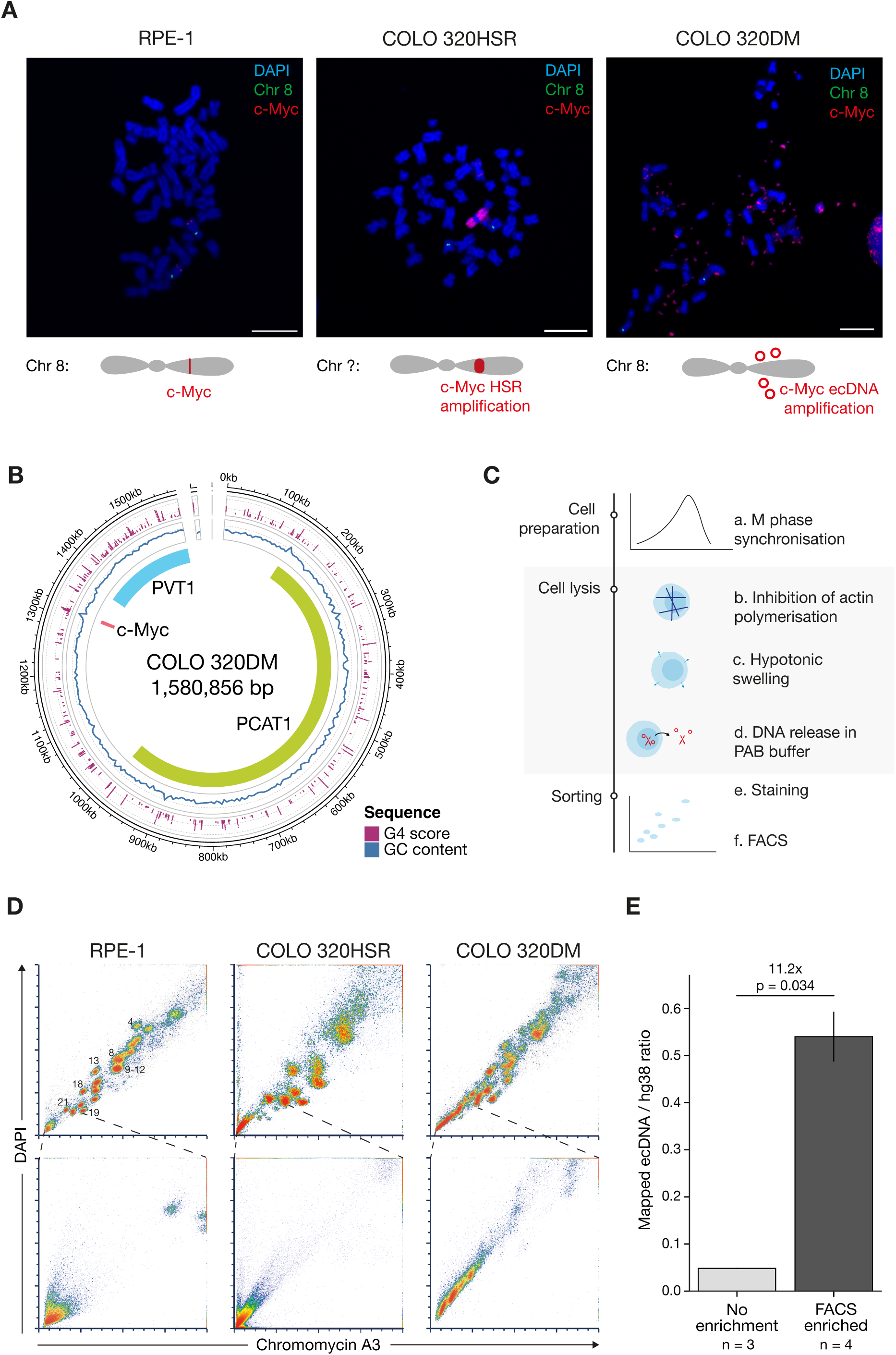
Characterisation of ecDNA in COLO 320DM and its isolation by FACS. **A.** Representative DNA FISH image of c-*MYC* localisation in RPE-1, COLO 320HSR and COLO 320DM. ecDNA or the HSR were labelled by FISH with a c-*MYC* probe. Slides were stained with DAPI, 5-fluorescein (centromeric region of Chromosome 8) and 5-TAMRA (c-*MYC*). Scale bar, 10 μm. The cartoons provide a graphic representation of the *MYC* amplification in COLO 320HSR and DM. **B.** Composite graph depicting the ecDNA structure in COLO 320DM generated using short reads and optical genome mapping with Amplicon Reconstructor. G4 calls from G4Hunter (36) above threshold 1.5 (purple dots); grey dotted lines are shown at G4Hunter scores 2.0 and 3.0); GC content (blue line); grey dotted line shown at 50%). **C.** Experimental workflow of the FACS-based protocol for ecDNA isolation involving cell preparation, DNA release and FACS sorting. **D.** Flow cytometry plots of chromosomes (top) and the region containing ecDNA and debris (bottom) from RPE-1 (left), COLO 320HSR (centre) and COLO 320DM (right) cell lines. The scales represent linear mean fluorescence intensity (MFI) but do not reflect detector voltage gains. Chromosome detection was performed using a detector gain of 622V (DAPI) and 651V (Chromomycin A3) for COLO 320DM; 622V (DAPI) and 604V (Chromomycin A3) for RPE-1; and 668V (DAPI) and 713V (Chromomycin A3) for COLO 320HSR. ecDNA detection was set using a gain of 750V (DAPI) and 750V (Chromomycin A3) for COLO 320DM; 750V (DAPI) and 750V (Chromomycin A3) for RPE-1; and 801V (DAPI) and 875V (Chromomycin A3) for COLO 320HSR. **E.** Ratio of ONT long read counts of sequencing reads aligned to the ecDNA region. Left: whole genome results excluding adaptive sampling sequencing, right: following FACS-based ecDNA purification. n indicates number of biological replicates. Error bars represent standard deviation.

**Figure 2.**
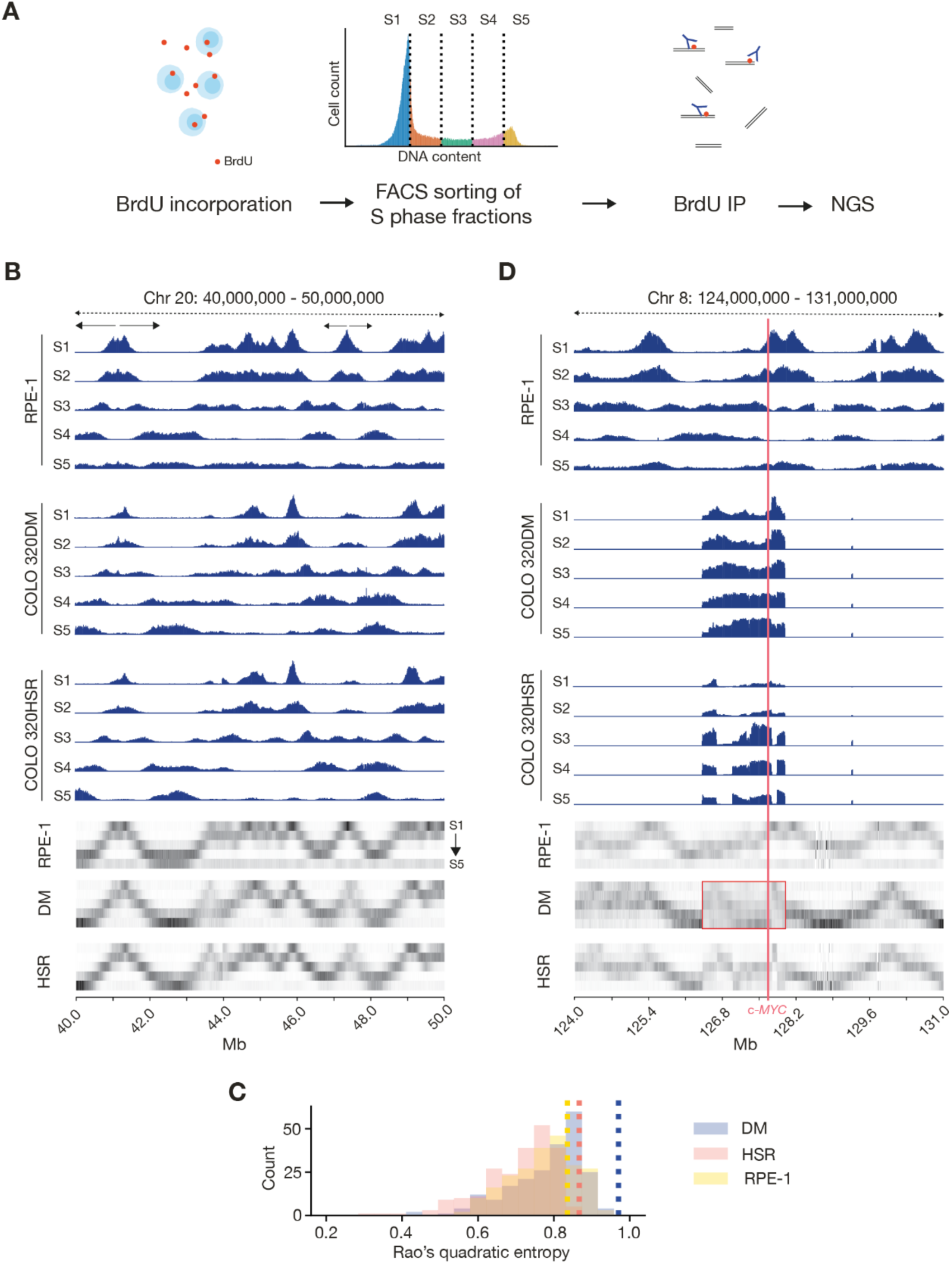
ecDNA replicates throughout S phase in COLO 320DM. **A.** Graphical representation of Repli-seq workflow. **B.** Replication timing analysis of a 10 Mb region on chromosome 20 (40,000,000–50,000,000) in RPE-1, COLO 320DM and COLO 320HSR. Coverage shown as reads per kilobase per million mapped reads (RPKM), group normalised to the highest peak in the visualised region between samples from the same cell line. Lower panels: visualisation of the proportion of reads in each S phase bin as a function of genome location. **C.** Rao entropy of replication timing. Replication timing was analysed using 1.6 Mb genomic bins (matching the ecDNA size in COLO 320DM), spaced 10 Mb apart. The Rao entropy for each bin was calculated for all three cell lines. Dashed lines indicate the Rao entropy of c-*MYC* gene in each respective cell line. **D.** Replication timing analysis of the region around ecDNA locus on chromosome 8 in RPE-1, COLO 320DM (DM) and COLO 320HSR (HSR). Coverage shown as RPKM, group normalised to the highest peak in the visualised region between samples from the same cell line. Lower panels: visualisation of the proportion of reads in each S phase bin as a function of genome location. ecDNA region highlighted with red box. c-*MYC* labelled in red. Note that the ecDNA regions in COLO 320DM and COLO 320HSR are shown aligned contiguously with adjacent sequence on chromosome 8 although the integration of the sequence in COLO 320HSR is not on chromosome 8 (Figure 1A).

Existing methods for ecDNA isolation pose significant challenges, requiring ecDNA cleavage or chromatin digestion, resulting in the extraction of limited data with high levels of noise (6, 39). To overcome these limitations, we refined established protocols for mitotic chromosome karyotyping by employing fluorescence-activated cell sorting (FACS) at the limits of its capabilities to detect and isolate ecDNA. In addition to previously tested approaches, we introduced Latrunculin B treatment to reduce the internal structural integrity of the cells by disrupting the actin cytoskeleton, which allowed us to omit the vortexing step. As a result, mitotic DNA, including both chromosomes and ecDNA, was released with minimal shearing, generating less debris and improving resolution, allowing for clearer visualisation of ecDNA that would otherwise be obscured by larger, chromosome-derived, DNA fragments (Figure 1C). This method is efficient in visualising and sorting intact chromosomes in chromosomally stable cell lines such as RPE-1 (Figure 1D), as well as in aneuploid cancer cell lines like COLO 320DM and COLO 320HSR (Figure 1D).

As anticipated, the substantial chromosomal heterogeneity observed in metaphase spreads of COLO 320DM led to less distinct chromosome profiles (Figure 1D). However, by increasing the voltage during sorting, we were able to detect low-molecular-weight DNA populations exclusive to ecDNA-positive cell lines, which were absent in ecDNA-negative COLO 320HSR and RPE-1 (Figure 1D), suggesting that these populations represent ecDNA rather than simple debris. Subsequent sequencing and copy number analysis revealed a 11 to 40-fold enrichment of ecDNA following FACS sorting compared to sequencing without prior isolation, further confirming that the isolated DNA molecules were ecDNA (Figure 1E).

### Replication Timing of ecDNA

To examine the timing of ecDNA replication, we employed the Repli-Seq technique, commonly used to assess replication timing in chromosomal DNA (22). We used five FACS gates across the cell cycle which provided the temporal resolution necessary not only to distinguish between early- and late-replicating regions, but also to assess whether ecDNA replication is synchronous within the population and whether it reflects the normal timing of the region when in a chromosomal context.

We performed Repli-Seq in COLO 320DM, COLO 320HSR and RPE-1. Each cell line was labelled with BrdU for 30 minutes, fixed in Carnoy’s fixative, stained with propidium iodide, and subjected to sorting into 5 separate fractions before deep sequencing of each fraction (Figure 2A).

To depict replication timing of ecDNA, we plotted the fraction of reads in windows of 10 kb across the genome for each S-phase bin (Figure 2B). Examination of a representative genomic region on chromosome 20 in the three cells lines suggested that they exhibit qualitatively similar replication timing profiles, with minor experiment-specific differences (Figure 2B; see Methods). To test this hypothesis across the genome, we measured replication synchronicity using the normalised mean Rao’s quadratic entropy (RQE) (30, 31), which is an information theory-based measure of distribution diversity, that considers the temporal ordering of the S-phase fractions (see Methods). In this analysis, a fragment replicating fully in a single fraction of S phase has a RQE of zero, while a fragment replicating uniformly across all five subphases achieves RQE of exactly 1. We plotted RQE of 1.6 Mb fragments (chosen to match the size of the ecDNA in COLO320DM) across the whole genome (Figure 2C). While differences in the distribution of synchronicity of replication timing are observed across the genome in the three lines, as would be expected for transformed cells (40, 41), there is significant overlap in the distributions. Notably, none of the 1.5Mb fragments sampled was found to replicate as asynchronously as ecDNA (Figure 2B).

However, replication of the ecDNA region in COLO 320DM appears considerably less synchronous than in the RPE-1 control with ongoing replication observed throughout the circle in the entire S phase (Figure 2D). This observation is supported by an RQE of 0.96, close to the maximum possible value of 1 and represents an outlier in the distribution of replication timing synchronicities in all three cell lines (Figure 2C). The same region in COLO 320HSR, in which the circle DNA is chromosomally integrated as an array, exhibits broadly late replication timing throughout, in contrast to the wave of replication that broadly passes left to right through the region in RPE-1 cells as S phase progresses (Figure 2D), but the overall synchronicity of the region is similar to that seen in RPE-1. Thus, replication on the ecDNA in COLO 320DM is significantly less ordered in terms of timing than the genome as a whole.

We next investigated replication fork velocity and origin distribution in the COLO320DM ecDNA.

### Replication origin distribution on ecDNA

Given the less synchronized replication observed in ecDNA (Figure 2), we explored whether ecDNA employs a different number and/or position of origins compared with the corresponding genomic sequence in a non-ecDNA context. We employed DNAscent (24) to identify replication forks and origins of replication in the sequences containing c-*MYC* amplified either on ecDNA or in the HSR. In brief, cells were incubated with two nucleoside analogues EdU and BrdU for 6 minutes each, which were incorporated into newly synthesised DNA during replication. The isolated DNA was then subjected to ultra-long ONT sequencing (Figure 3A). The characteristic disruptions in electrical current caused by EdU and BrdU as the labelled DNA passes through the sequencing pores allows the probability of their presence to be calculated by the DNAscent model on top of the canonical DNA sequence. The distribution of EdU and BrdU along the individual DNA molecules is indicated as a probability (Figure 3B). Applying density-based segmentation to these probabilities creates replication tracts that identify fork direction, velocity, stalling and replication origins (24).

**Figure 3:**
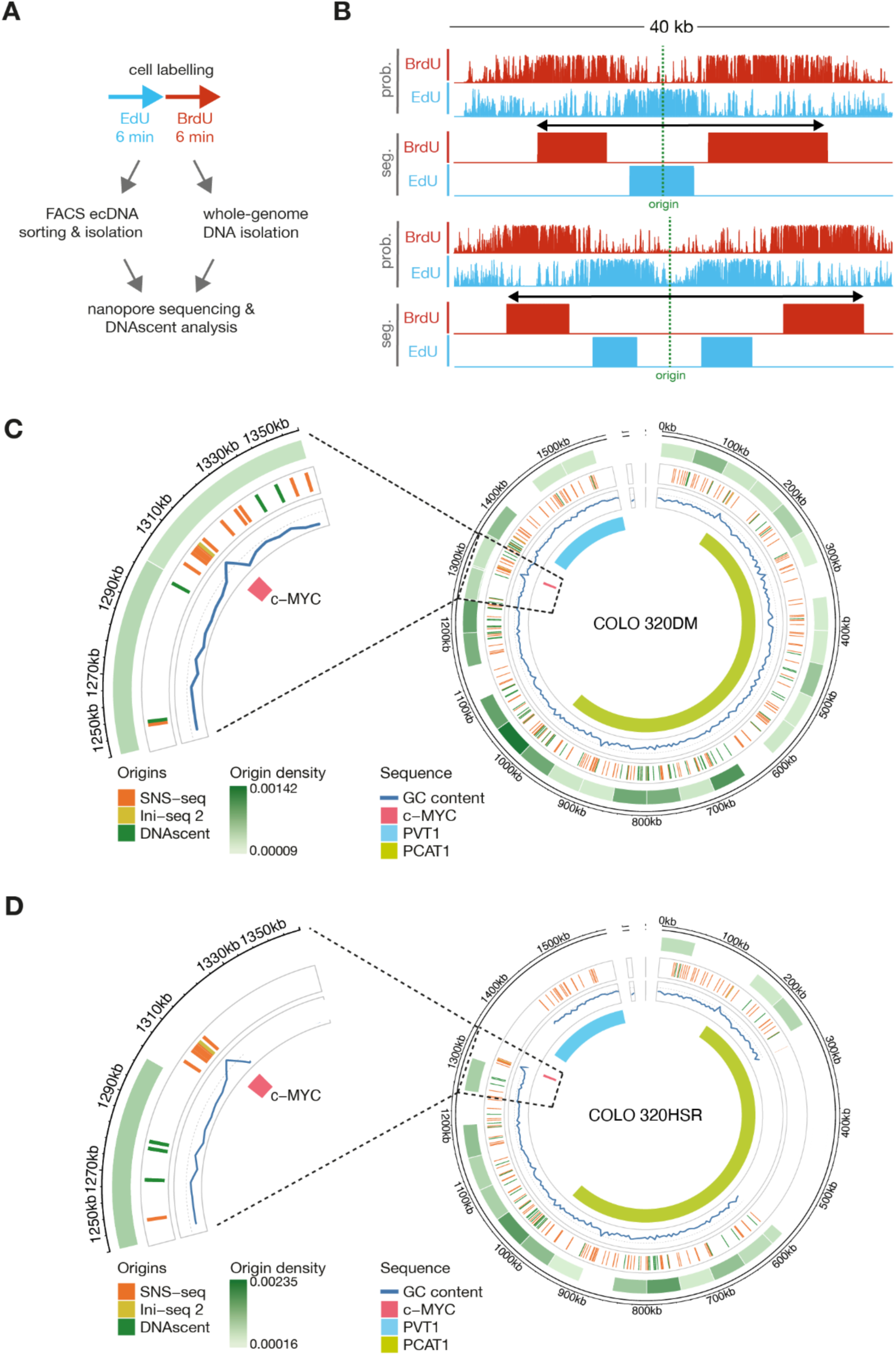
Location of origins of replication. **A.** Schematic of base analogue pulsing protocol followed by either FACS sorting or whole genome DNA extraction for subsequent ultra-long Nanopore sequencing and DNAscent analysis. **B.** Representative raw base analogue incorporation probabilities and DNAscent segmentation of origins of replication each represented by four tracks (Upper tracks: Raw BrdU and EdU probabilities (prob.) at thymidine positions; Lower tracks BrdU and EdU segmentation (seg.) derived from the raw probabilities). Both origins map to ecDNA in the untreated COLO 320DM cell line. (Coordinates on ecDNA map to chromosome chr8_126425747-127997820 bp. The top origin track maps within the reconstructed ecDNA between 793,008 to 801,870 bp; the lower example to 470,941 to 470,941 bp). Raw DNAscent probabilities for EdU or BrdU are shown on a scale from 0 to 1, segmentation is binary. **C. & D.** Distribution of origins in C. COLO 320DM and D. Colo 320HSR on the ecDNA reference map (Low density = light green shading to high density = dark green shading). Orange: SNS-seq origin locations (43); yellow: Ini-seq 2 origins (44); green: DNAscent origin locations; Blue line: GC content. Zoom in shows the region around c-*MYC* (1250 kb to 1350 kb).

We initially employed DNAscent on FACS-isolated ecDNA. While this allowed identification of replication forks and replication origins in ecDNA (Supplementary Figure 1), the throughput was not satisfactory and the need to concentrate the circle DNA from the relatively large volumes generated by FACS sorting led to DNA breakage and relatively short reads in the ONT sequencing runs (Supplementary Table 1). We therefore moved to using whole genome sequencing using the ONT PromethION instrument.

Using PromethION sequencing on unselected DNA we were able to achieve an average N50 of 91.1 +/-16.7 kb (Supplementary Table 1). From this data, we identified 1312 reads with origin of replication calls in COLO 320DM and 1115 reads with origin calls in COLO 320HSR DNA. Of these, 81 and 43 respectively mapped to the 1.6 Mb interval covered by the ecDNA (Supplementary Table 1). The sites to which origins were mapped in the ecDNA interval exhibited no significant increase in GC content or G4 prevalence over randomly selected regions from the same interval (Supplementary Figure 2 & 3). We examined whether the distribution of origins is conserved between these amplification types, given differences in replication timing (Figure 2D). There is not a significantly higher origin density in the ecDNA of COLO 320DM compared with the equivalent interval in COLO 320HSR, consistent with the genome-wide analysis (Supplementary Figure 4). Further specific analysis of the region around c-*MYC*, which contains known sites of replication initiation (42), revealed origins near the c-*MYC* and *PVT-1* loci in both cell lines. However, these did not align with the origins within c-*MYC* gene observed in other systems (Figure 3C & D), highlighting potential differences in the replication patterns of the amplified c-*MYC* locus in COLO320 cell lines (43, 44) and consistent with the disorganised replication timing of this region in the COLO320 DM ecDNA (Figure 2). Origin distribution across the ecDNA differed significantly from a uniform distribution based on origin counts in 50kb segments for all 3 conditions as determined by a one-sample Kolmogorov-Smirnov test (COLO 320DM: KS statistic = 0.91, p-value = 1.9e-34; COLO 320HSR KS statistic = 0.76, p-value = 7.6e-9; COLO320 DM HU-treated KS statistic = 0.92, p-value = 5.9e-36).

### DNA replication is slower on ecDNA compared to chromosomal DNA

To further investigate the differences in replication dynamics between ecDNA and the chromosomes, we measured replication fork velocity and stalling in ecDNA-positive and ecDNA-negative cell lines. Using ONT sequencing and DNAscent, we detected individual nascent DNA fragments labelled with EdU and BrdU (Figure 4A). By dividing the length of continuous stretches of labelled DNA by the pulse durations we calculated replication fork velocity - longer tracts indicate faster overall fork progression. Additionally, we examined the pattern of signal decay to assess fork stalling: an abrupt loss of signal corresponds to replication fork stalling or termination, while a gradual decline reflects normal fork progression as BrdU levels diminish over time due to the thymidine chase (Figure 4A). This approach enabled us to directly measure and compare replication fork dynamics on ecDNA and chromosomal DNA, revealing statistically significant differences in fork velocity and stalling patterns (24).

**Figure 4.**
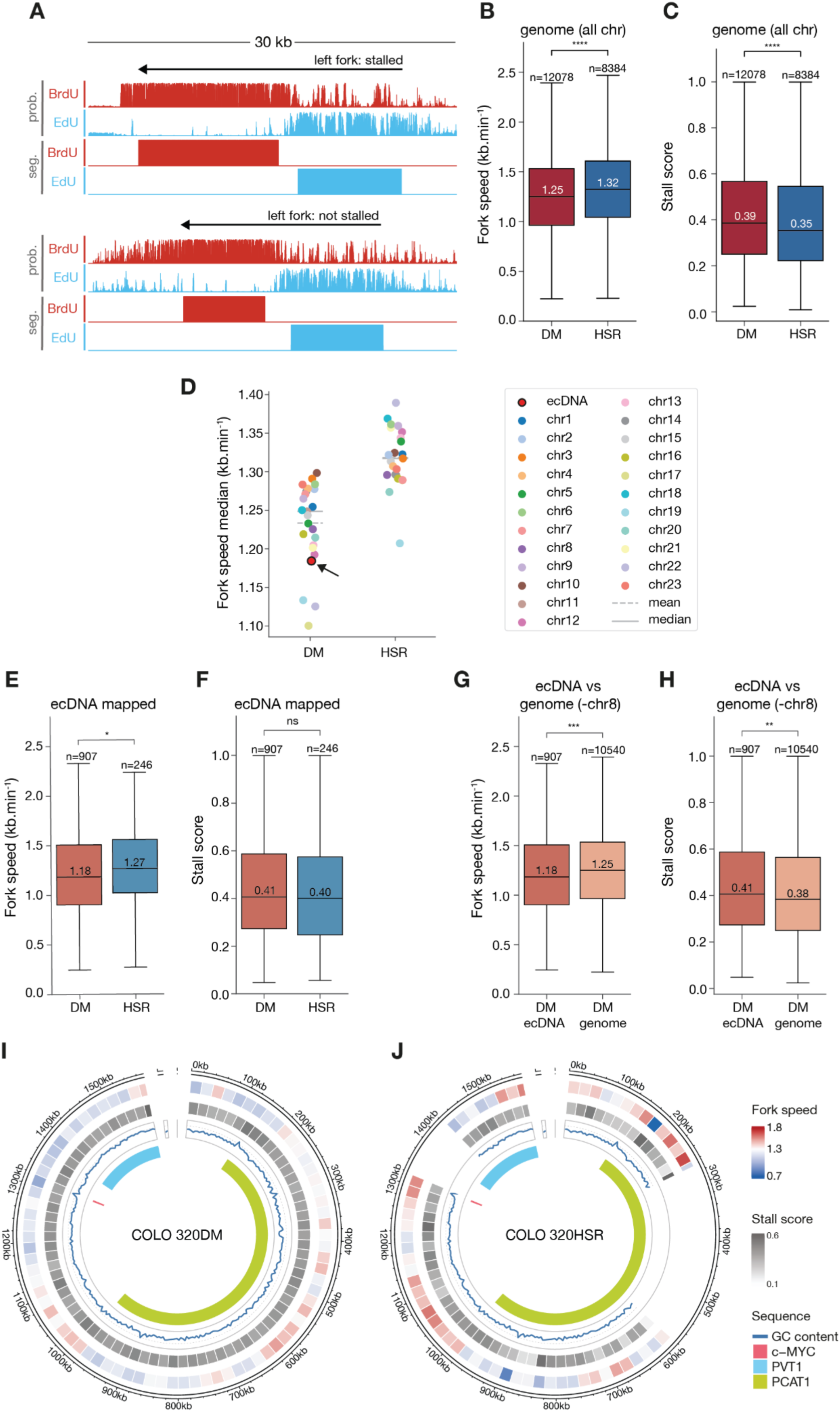
Replication dynamics on ecDNA. **A.** Representative DNAscent tracks for leftward-moving replication forks (BrdU probability (prob.) and segmentation (seg): red; EdU: blue. The top example shows a stalled fork, characterised by a sharp drop-off in BrdU incorporation probability at the fork tip (upper tracks; stall score of 0.9991, fork speed of 1.79 kb/min). The lower example tracks show a replication fork with no fork stalling (stall score of 0.2657, fork speed 1.36 kb/min) represented by a smooth decrease in BrdU probabilities at the fork tip. Both examples are from the Colo 320HSR cell line within the ecDNA region from chromosome 8. Raw DNAscent probabilities for EdU or BrdU are shown on a scale from 0 to 1, segmentation is binary. **B.** & **C.** Comparison of genome-wide fork speeds (B) and stall scores (C) in COLO 320DM (DM; red) and Colo 320HSR (HSR; blue). **D.** Median fork speed per chromosome in COLO 320DM (DM) and COLO 320HSR (HSR) cell lines. For the COLO 320DM cell line, the median fork speed on the ecDNA is shown as a red dot with black outline. **E.** & **F.** Comparison of fork speed (E) and stall scores (F) of forks mapped to the ecDNA interval in COLO 320DM (circular, light red) and COLO 320HSR (chromosomally reintegrated, light blue). **G.** & **H.** Comparison of fork speed (G) and stall scores (H) within COLO 320DM cell line between forks mapped to the ecDNA interval (dark orange) and forks mapped to chromosomes (excluding chromosome 8 from which the ecDNA originates, light orange). **I.** & **J.** Visualisation of replication fork speeds (I) and stall scores (J) averaged across 20kb segments of the ecDNA interval in COLO 320DM and **J.** COLO 320HSR cell line. Outer track, fork speeds; second track, stall score; third track, GC content (blue line); Inner track, genes. Fork speeds and stall scores representation uses the same range in I and J with the most extreme values across both cell lines determining the minimum and maximum colour shades. All p-values for boxplots with a fork speed are obtained from a two-sided Welch’s t-test with no assumption of equal variances and all p-values in boxplots for stall scores are obtained from a two-sided non-parametric Wilcoxon Rank Sum test. Statistical significance: ns = not significant (p ≥ 0.05), * p < 0.05, ** p < 0.01, *** p < 0.001, **** p < 0.0001.

Global fork velocity was reduced genome-wide in ecDNA-positive COLO 320DM compared to ecDNA-null COLO 320HSR (Figure 4B). Moreover, ecDNA-positive cells displayed a higher frequency of fork stalling (Figure 4C), though this increase in stalling did not account for the reduced fork velocity since forks selected for matched levels of stalling still exhibited reduced fork velocity in COLO 320DM (Supplementary Figure 5). Consistent with these findings, replication forks on each chromosome in COLO 320HSR were progressing more rapidly than on its counterpart in COLO 320DM cells (Figure 4D).

To further characterise ecDNA replication, we compared the replication dynamics of ecDNA with its corresponding amplified sequence on the HSR in COLO 320HSR cells. Replication of the ecDNA was approximately 6.5% slower (1.27 kb/min vs 1.18 kb/min), although no significant difference in fork stalling rates was observed (Figure 4E & F). Similarly, ecDNA replication in COLO 320DM was slower than chromosomal DNA replication (1.18 kb/min vs 1.25 kb/min; Figure 4G & H), with a marginally higher stalling rate (0.41 vs 0.38).

We next examined the distribution of replication fork velocities across COLO 320DM ecDNA and the corresponding sequence in COLO 320HSR. Mean fork velocity, visualised in 20 kb bins, ranged from 0.7 to 1.8 kb/min, with most regions showing concordant replication velocities between ecDNA and HSR (Figure 4I & J). However, regions such as the c-*MYC* locus exhibited stark differences in replication dynamics. While the c-*MYC* region was one of the fastest-replicating in the HSR, it was the slowest-replicating on ecDNA, indicating substantial differences in replication dynamics between ecDNA and chromosomal DNA (Figure 4I & J). A similar pattern was observed for the stall rate (Figure 4I & J), with ecDNA displaying elevated stalling in regions where HSR replication was most efficient. These findings suggest that ecDNA replication diverges significantly from chromosomal HSR replication, even in the regions with the same underlying sequence and, overall, consistent with replication on ecDNA being more intrinsically stressed (17).

### Hydroxyurea disrupts ecDNA replication, leading to its depletion from cells

Consistent with previous studies (18–21), hydroxyurea treatment led to a marked reduction in ecDNA levels in COLO 320DM cells (Figure 5A). This was reversed on withdrawing HU (Figure 5A), suggesting strong selective pressure to maintain ecDNA in these cells. To further explore this phenomenon, we analysed ecDNA replication dynamics under HU treatment using DNAscent.

**Figure 5.**
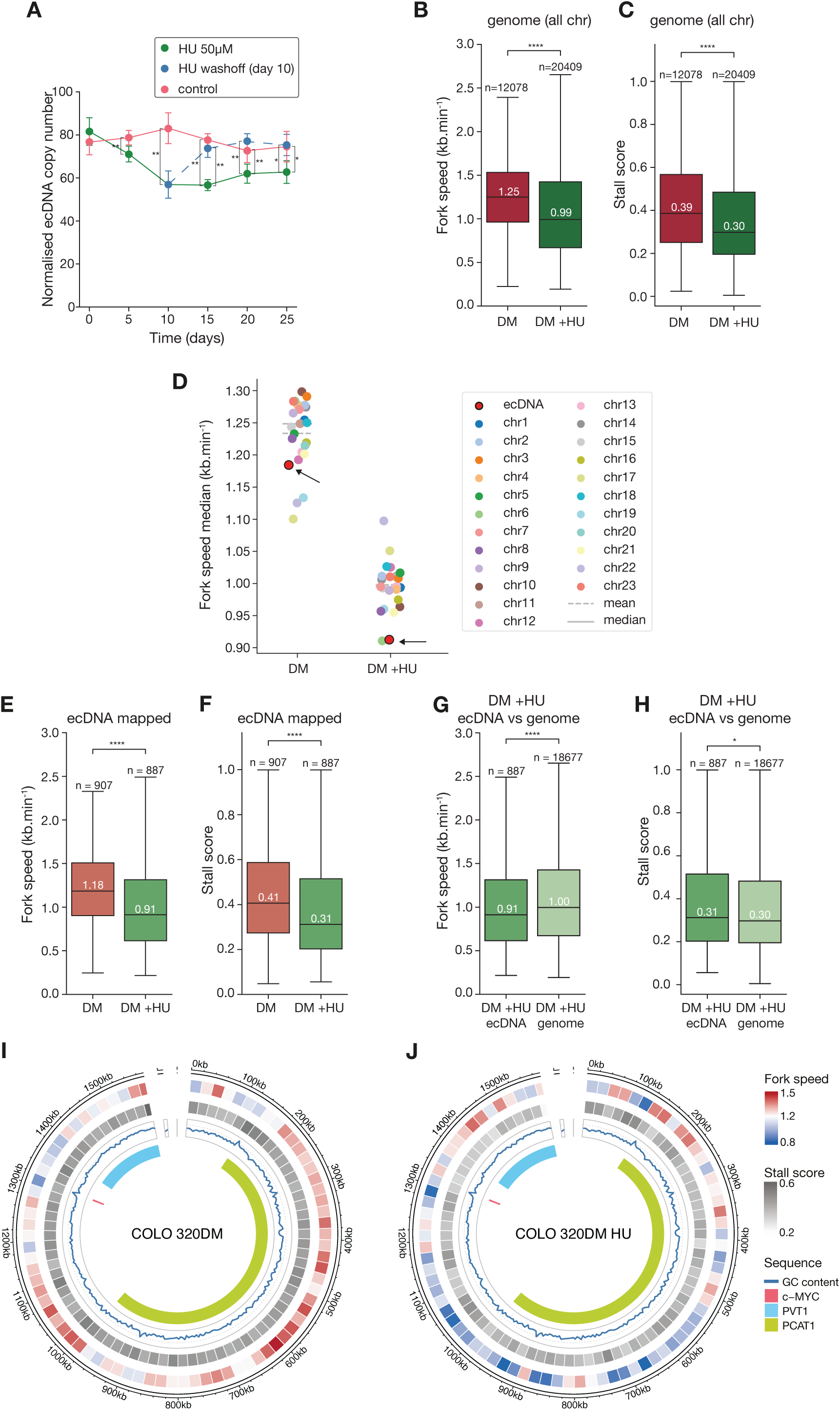
Hydroxyurea treatment slows down DNA replication on ecDNA. **A.** Copy number variation, estimated by qPCR, in COLO 320DM with HU treatment (green), upon HU removal (blue) and without treatment (red). Pair-wise comparisons are assessed with a two-sided Wilcoxon Rank Sum test. **B.** & **C.** Fork speeds (C) and stall scores (D) in untreated (red) and HU-treated (green) COLO 320DM cell lines. **D.** Median fork speed per chromosome in untreated (DM) and HU treated (DM_HU) COLO 320DM cell lines. The median fork speed on ecDNA is shown as a red dot with a black edge. **E.** & **F.** Comparison of fork speed (F) and stall scores (G) stall on ecDNA region mapped forks in untreated (medium red) and HU-treated (medium green) COLO 320DM cell line. **G.** & **H.** Comparison of fork speed (H) and stall scores (I) in forks mapped to ecDNA (medium green) and forks mapped to chromosomes (excluding chromosome 8), light green from HU-treated COLO 320DM cells. **I.** & **J.** Visualisation of replication fork speeds and stall scores averaged across 20kb segments in the ecDNA interval of untreated (J) and HU-treated (K) COLO 320DM cells. Fork speeds lower than the average fork speed (light grey) are shaded in blue, higher in red and stall score averages are all low to moderate (light to medium grey). GC content is shown as a blue line and position of c-*MYC* and *PVT1* genes are highlighted by pink and blue blocks, respectively. Colouring of fork speeds and stall scores is the same in I and J with the most extreme values across both cell lines setting the minimum and maximum colour shades. All p values for panel A are obtained from a Wilcoxon Rank Sum test. All p-values for boxplots with a fork speed are obtained from a two-sided Welch’s t-test with no assumption of equal variances and all p-values in boxplots for stall scores are obtained from a two-sided non-parametric Wilcoxon Rank Sum test. Statistical significance: ns = not significant (p ≥ 0.05), * p < 0.05, ** p < 0.01, *** p < 0.001, **** p < 0.0001.

First, we investigated changes in origin distribution under HU treatment, as reduced fork velocity can trigger activation of dormant origins to compensate for slower replication (45, 46). Upon HU treatment, we observed increased origin activation both genome-wide and on ecDNA (Supplementary Figure 4), particularly around the c-*MYC* locus (Supplementary Figure 2). This shows that dormant origins are activated both genome-wide and on ecDNA under replication stress.

Hydroxyurea treatment also resulted in a genome-wide reduction in fork velocity of 20.7% (Figure 5B; 0.99 vs 1.25 kb/min), with every chromosome exhibiting decreased replication speed (Figure 5D). Notably, DNAscent-detected fork stalling also decreased in HU treated cells (Figure 5C). This suggests that the reduced velocity of DNA synthesis resulting from nucleotide depletion leads to fewer sharply cut-off tracts of DNA synthesis, which likely represent stalled forks (24), is perhaps counterintuitive. However, such resolution of stalls has previous not been accessible to techniques like DNA combing due to their inability to distinguish between gradual decline of the signal, and abrupt signal loss suggesting stalling (47). The observed reduction in the stall score may reflect either the slower forks in HU less frequently encountering synthesis impediments, or the slower replication allows more opportunity to deal with problems on the template ‘on-the-fly’ allowing synthesis to continue. This reduction in fork velocity, observed both on ecDNA and genome-wide, may also lead to underreplication, which might be mitigated by the increased activation of dormant origins in response to HU-induced replication stress.

When comparing specifically ecDNA replication under HU treatment, the replication slowing effect was even more pronounced. Fork velocity on ecDNA dropped significantly by 22.9% (0.91 vs 1.18 kb/min), making it the slowest replicating fraction relative to all chromosomes (Figure 5D & E). The stalling rate also decreased from 0.41 to 0.31 (Figure 5F), indicating that stalled forks do not account for the overall reduction in replication speed. Even under HU treatment, ecDNA replication remained slower than chromosomal DNA (Figure 5G; 0.91 vs 1 kb/min), with an even higher relative decrease in replication velocity (5.6% vs 9% reduction) (Supplementary Figure 7). Notably, the stalling rate across both fractions dropped to nearly identical levels (0.31 vs 0.30) (Figure 5H). These effects were observed in both ecDNA and chromosomal DNA but to different extents. Replication fork velocity decreased proportionally more on ecDNA, while the stalling rate was similarly reduced in both fractions (Supplementary Figure 7), which may contribute to the loss of ecDNA from cells under HU treatment.

Further analysis of 20 kb fragments across ecDNA confirmed a significant reduction in both replication velocity and stalling (Figure 5I & J). However, the distribution of fast- and slow-replicating regions shifted markedly. Regions surrounding c-*MYC* and PVT-1, previously among the slowest replicating areas, became the fastest replicating upon HU treatment, even surpassing their pre-treatment speeds. This suggests that HU has its greatest impact on the fastest replicating regions, indicating a complex, region-specific influence on ecDNA replication. In contrast, the stalling rate across the other regions of ecDNA decreased more uniformly, suggesting a differential impact of HU on replication dynamics that may be driven by the variations in underlying sequence or local chromatin environment (Figure 5I & J).

## Discussion

This study reveals that ecDNA replicates asynchronously throughout the S phase, contrasting with the well-defined replication timing of linear chromosomal DNA. This asynchrony suggests that ecDNA lacks the strict temporal regulation of replication initiation found in chromosomal DNA, possibly due to its previously reported generally accessible euchromatic nature (7, 48).

Given this largely asynchronous pattern of replication on ecDNA, we explored whether ecDNA employs a different number or arrangement of replication origins, compared with the corresponding chromosomal sequence. This property of ecDNA replication could influence both replication dynamics and potential replication-transcription conflicts. The analysis of the distribution of replication origins revealed an increased number of origins on ecDNA upon HU treatment. The activation of additional origins on ecDNA under replication stress, suggests that the well-documented compensatory mechanism by which cryptic origins are fired to ensure complete replication despite slower fork progression (45, 46) is active on ecDNA. This increased origin firing may also reflect the less strict replication pattern of ecDNA, act as an adaptation to increased replication stress and may also contribute to replication–transcription conflicts, given the high transcriptional activity of oncogenes on ecDNA (2, 7, 17).

The mildly reduced fork velocity and elevated stalling rates on ecDNA under normal conditions suggest that replication forks on ecDNA are constitutively subject to some degree of replication stress. Hydroxyurea (HU) treatment results in an expected significant decrease in fork speed, but surprisingly also causes a reduction in stalling, suggesting that slower replication forks are less prone to complete stalling of DNA synthesis. This finding contrasts with previous reports that HU treatment is linked to increased fork stalling (49). One explanation is linked to the fact that previously used DNA fibre combing analyses could not distinguish between slower replication and increased stalling. In contrast, our observations reveal that the reduced fork velocity reflects genuine slowing of the replication fork rather than increased stalling, as DNAscent enables the differentiation of a gradual decay of the signal - suggesting slower progression - and a sudden stop, suggesting fork stalling. This distinction cannot be achieved using traditional DNA fibre methods.

We also hypothesise that the depletion of ecDNA under HU treatment is likely due to its vulnerability to replication stress, as suggested by more marked slowing of replication on ecDNA compared with chromosomal DNA in the presence of HU. This supports the potential therapeutic strategy of inducing replication stress to selectively target ecDNA-bearing tumour cells (11, 17, 19).

Applying DNAscent to ecDNA allowed us to map replication origins with high precision on single DNA molecules without requiring amplification, reducing noise and artefacts associated with other high-throughput methods. Our study offers new insights into the distinct replication dynamics of ecDNA. In addition, we developed a new method for ecDNA isolation, FINE, which minimises ecDNA processing and offers high specificity, potentially enabling the study of ecDNA properties such as chromatin composition and three-dimensional structure through proteomics and imaging. While FINE is limited by relatively low yields of ecDNA in large volumes and incompatibility with some replication assays like Repli-seq, it provides a potentially valuable tool for ecDNA detection and characterisation, e.g. through proteomics or structural analysis.

While our study sheds light on ecDNA replication dynamics, it raises questions about the molecular mechanisms governing replication initiation and regulation on ecDNA. The roles of specific proteins or epigenetic marks in the altered replication timing and origin activation patterns remain to be elucidated. Notably, ecDNA exhibits a more open chromatin conformation than chromosomal DNA (7), which may facilitate replication by improving the accessibility to the replication machinery, or conversely, impede it by generating high levels of replication-transcription conflicts (17). Furthermore, the mechanisms governing replication licensing on ecDNA remains unresolved. If, consistent with the previous reports (16), ecDNA is replicated only once per cell cycle, its copy number can increase only through random segregation and subsequent selection, rather than through multiple rounds of replication (50). In any case, understanding ecDNA licensing remains a fundamental issue that requires further investigation. Additionally, the interplay between replication and transcription machinery on ecDNA warrants further investigation, particularly given the potential for replication– transcription conflicts to contribute to genomic instability (17, 51). We hypothesise that increased transcription (7, 17) together with slower ecDNA replication (particularly under HU treatment), and the euchromatic state of ecDNA may collectively contribute to the dysregulated origin landscape.

Finally, ecDNA biology exhibits radical heterogeneity between cell lines and even among individual ecDNA molecules. Therefore, future studies should examine the applicability of these findings across cancer types and ecDNA compositions. Understanding whether the vulnerabilities of ecDNA replication can be generalized will be crucial for developing targeted therapeutic interventions. The application of advanced replication analysis techniques and improved ecDNA isolation methods opens new avenues for research into diverse aspects of ecDNA biology and its role in cancer progression. Investigating how cancer cells compensate for ecDNA loss and whether they can develop resistance to replication-targeting therapies will determine future treatment strategies.

## Data Availability

Sequencing data can be accessed at the Gene Expression Omnibus archive with the accession number GSE186675. The code for the ini-seq 2 origin caller can be found at https://github.com/Sale-lab. The ONT sequencing data can be found on ENA under the accession number PRJEB83636. The code for the DNAscent-associated analyses is publicly available under https://github.com/Pfuderer/ecDNA_replication_dynamics.

## Acknowledgements

We would like to thank the FACS facility at the MRC Laboratory of Molecular Biology, especially P. A. Penttilä, M. Daly, Y. Li, F. Zhang, and D. Nolan, for support. We also thank Alastair Crisp for support with AmpliconArchitect ecDNA structure analysis, Welcome Sanger Sequencing R&D facility, in particular Iraad Bronner for Nanopore and OGM sequencing, and members of the Sale and Chin labs for helpful discussions.

## Contributions

J.J.J. and J.E.S. conceptualised the project. J.J.J., G.P., P.L.P., J.E.S., and M.A.B. designed the experiments. J.J.J. performed wet lab experiments and conducted bioinformatic Repli-seq analyses. P.L.P. carried out bioinformatic analyses of DNA replication fork dynamics. P.C. performed replication timing statistical analyses and visualisation. J.J.J. and P.L.P. wrote the manuscript with input from all authors. J.E.S. and M.A.B. supervised the project.

## References

1. Storlazzi, C.T., Lonoce, A., Guastadisegni, M.C., Trombetta, D., D’Addabbo, P., Daniele, G., L’Abbate, A., Macchia, G., Surace, C., Kok, K., Ullmann, R., Purgato, S., Palumbo, O., Carella, M., Ambros, P.F. and Rocchi, M. (2010) Gene amplification as double minutes or homogeneously staining regions in solid tumors: origin and structure. Genome Res 20, 1198–1206.

2. Turner, K.M., Deshpande, V., Beyter, D., Koga, T., Rusert, J., Lee, C., Li, B., Arden, K., Ren, B., Nathanson, D.A., Kornblum, H.I., Taylor, M.D., Kaushal, S., Cavenee, W.K., Wechsler-Reya, R., Furnari, F.B., Vandenberg, S.R., Rao, P.N., Wahl, G.M., Bafna, V. and Mischel, P.S. (2017) Extrachromosomal oncogene amplification drives tumour evolution and genetic heterogeneity. Nature 543, 122–125.

3. Verhaak, R.G.W., Bafna, V. and Mischel, P.S. (2019) Extrachromosomal oncogene amplification in tumour pathogenesis and evolution. Nat Rev Cancer 19, 283–288.

4. Bailey, C., Pich, O., Thol, K., Watkins, T.B.K., Luebeck, J., Rowan, A., Stavrou, G., Weiser, N.E., Dameracharla, B., Bentham, R., Lu, W.-T., Kittel, J., Yang, S.Y.C., Howitt, B.E., Sharma, N., Litovchenko, M., Salgado, R., Hung, K.L., Cornish, A.J., Moore, D.A., Houlston, R.S., Bafna, V., Chang, H.Y., Nik-Zainal, S., Kanu, N., McGranahan, N., Genomics England Consortium, Flanagan, A.M., Mischel, P.S., Jamal-Hanjani, M. and Swanton, C. (2024) Origins and impact of extrachromosomal DNA. Nature 635, 193–200.

5. Kim, H., Nguyen, N.P., Turner, K., Wu, S., Gujar, A.D., Luebeck, J., Liu, J., Deshpande, V., Rajkumar, U., Namburi, S., Amin, S.B., Yi, E., Menghi, F., Schulte, J.H., Henssen, A.G., Chang, H.Y., Beck, C.R., Mischel, P.S., Bafna, V. and Verhaak, R.G.W. (2020) Extrachromosomal DNA is associated with oncogene amplification and poor outcome across multiple cancers. Nat Genet 52, 891–897.

6. Koche, R.P., Rodriguez-Fos, E., Helmsauer, K., Burkert, M., MacArthur, I.C., Maag, J., Chamorro, R., Munoz-Perez, N., Puiggròs, M., Dorado Garcia, H., Bei, Y., Röefzaad, C., Bardinet, V., Szymansky, A., Winkler, A., Thole, T., Timme, N., Kasack, K., Fuchs, S., Klironomos, F., Thiessen, N., Blanc, E., Schmelz, K., Künkele, A., Hundsdörfer, P., Rosswog, C., Theissen, J., Beule, D., Deubzer, H., Sauer, S., Toedling, J., Fischer, M., Hertwig, F., Schwarz, R.F., Eggert, A., Torrents, D., Schulte, J.H. and Henssen, A.G. (2020) Extrachromosomal circular DNA drives oncogenic genome remodeling in neuroblastoma. Nat Genet 52, 29–34.

7. Wu, S., Turner, K.M., Nguyen, N., Raviram, R., Erb, M., Santini, J., Luebeck, J., Rajkumar, U., Diao, Y., Li, B., Zhang, W., Jameson, N., Corces, M.R., Granja, J.M., Chen, X., Coruh, C., Abnousi, A., Houston, J., Ye, Z., Hu, R., Yu, M., Kim, H., Law, J.A., Verhaak, R.G.W., Hu, M., Furnari, F.B., Chang, H.Y., Ren, B., Bafna, V. and Mischel, P.S. (2019) Circular ecDNA promotes accessible chromatin and high oncogene expression. Nature 575, 699–703.

8. Hung, K.L., Jones, M.G., Wong, I.T.-L., Curtis, E.J., Lange, J.T., He, B.J., Luebeck, J., Schmargon, R., Scanu, E., Brückner, L., Yan, X., Li, R., Gnanasekar, A., Chamorro González, R., Belk, J.A., Liu, Z., Melillo, B., Bafna, V., Dörr, J.R., Werner, B., Huang, W., Cravatt, B.F., Henssen, A.G., Mischel, P.S. and Chang, H.Y. (2024) Coordinated inheritance of extrachromosomal DNAs in cancer cells. Nature 635, 201–209.

9. Lange, J.T., Rose, J.C., Chen, C.Y., Pichugin, Y., Xie, L., Tang, J., Hung, K.L., Yost, K.E., Shi, Q., Erb, M.L., Rajkumar, U., Wu, S., Taschner-Mandl, S., Bernkopf, M., Swanton, C., Liu, Z., Huang, W., Chang, H.Y., Bafna, V., Henssen, A.G., Werner, B. and Mischel, P.S. (2022) The evolutionary dynamics of extrachromosomal DNA in human cancers. Nat Genet 54, 1527–1533.

10. Shoshani, O., Brunner, S.F., Yaeger, R., Ly, P., Nechemia-Arbely, Y., Kim, D.H., Fang, R., Castillon, G.A., Yu, M., Li, J.S.Z., Sun, Y., Ellisman, M.H., Ren, B., Campbell, P.J. and Cleveland, D.W. (2021) Chromothripsis drives the evolution of gene amplification in cancer. Nature 591, 137–141.

11. Nathanson, D.A., Gini, B., Mottahedeh, J., Visnyei, K., Koga, T., Gomez, G., Eskin, A., Hwang, K., Wang, J., Masui, K., Paucar, A., Yang, H., Ohashi, M., Zhu, S., Wykosky, J., Reed, R., Nelson, S.F., Cloughesy, T.F., James, C.D., Rao, P.N., Kornblum, H.I., Heath, J.R., Cavenee, W.K., Furnari, F.B. and Mischel, P.S. (2014) Targeted therapy resistance mediated by dynamic regulation of extrachromosomal mutant EGFR DNA. Science 343, 72–76.

12. Deng, X., Zhang, L., Zhang, Y., Yan, Y., Xu, Z., Dong, S. and Fu, S. (2006) Double minute chromosomes in mouse methotrexate-resistant cells studied by atomic force microscopy. Biochem Biophys Res Commun 346, 1228–1233.

13. Hahn, P.J., Nevaldine, B. and Longo, J.A. (1992) Molecular structure and evolution of double-minute chromosomes in methotrexate-resistant cultured mouse cells. Mol Cell Biol 12, 2911–2918.

14. Kaufman, R.J., Brown, P.C. and Schimke, R.T. (1979) Amplified dihydrofolate reductase genes in unstably methotrexate-resistant cells are associated with double minute chromosomes. Proceedings of the National Academy of Sciences 76, 5669–5673.

15. Levan, A. and Levan, G. (1978) Have double minutes functioning centromeres? Hereditas 88, 81–92.

16. Barker, P.E., Drwinga, H.L., Hittelman, W.N. and Maddox, A.M. (1980) Double minutes replicate once during S phase of the cell cycle. Exp Cell Res 130, 353–360.

17. Tang, J., Weiser, N.E., Wang, G., Chowdhry, S., Curtis, E.J., Zhao, Y., Wong, I.T.-L., Marinov, G.K., Li, R., Hanoian, P., Tse, E., Mojica, S.G., Hansen, R., Plum, J., Steffy, A., Milutinovic, S., Meyer, S.T., Luebeck, J., Wang, Y., Zhang, S., Altemose, N., Curtis, C., Greenleaf, W.J., Bafna, V., Benkovic, S.J., Pinkerton, A.B., Kasibhatla, S., Hassig, C.A., Mischel, P.S. and Chang, H.Y. (2024) Enhancing transcription-replication conflict targets ecDNA-positive cancers. Nature 635, 210–218.

18. Shimizu, N., Nakamura, H., Kadota, T., Kitajima, K., Oda, T., Hirano, T. and Utiyama, H. (1994) Loss of amplified c-myc genes in the spontaneously differentiated HL-60 cells Cancer Research 54, 3561–3567.

19. 19. Von Hoff, D.D., McGill, J.R., Forseth, B.J., Davidson, K.K., Bradley, T.P.R.V.D.D., Van Devanter, D.R. and Wahl, G.M. (1992) Elimination of extrachromosomally amplified MYC genes from human tumor cells reduces their tumorigenicity. Proceedings of the National Academy of Sciences 89, 8165–8169.

20. Von Hoff, D.D., Waddelow, T., Forseth, B., Davidson, K., Scott, J. and Wahl, G. (1991) Hydroxyurea accelerates loss of extrachromosomally amplified genes from tumor cells Cancer research 51, 6273–6279.

21. 21. Eckhardt, S.G., Dai, A., Davidson, K.K., Forseth, B.J., Wahl, G.M. and Von Hoff, D.D. (1994) Induction of differentiation in HL60 cells by the reduction of extrachromosomally amplified c-myc. Proceedings of the National Academy of Sciences 91, 6674–6678.

22. Zhao, P.A., Sasaki, T. and Gilbert, D.M. (2020) High-resolution Repli-Seq defines the temporal choreography of initiation, elongation and termination of replication in mammalian cells. Genome Biol 21, 76.

23. Marchal, C., Sasaki, T., Vera, D., Wilson, K., Sima, J., Rivera-Mulia, J.C., Trevilla-García, C., Nogues, C., Nafie, E. and Gilbert, D.M. (2018) Genome-wide analysis of replication timing by next-generation sequencing with E/L Repli-seq. Nat Protoc 13, 819–839.

24. Boemo, M.A. (2021) DNAscent v2: detecting replication forks in nanopore sequencing data with deep learning. BMC Genomics 22, 430.

25. Schindelin, J., Arganda-Carreras, I., Frise, E., Kaynig, V., Longair, M., Pietzsch, T., Preibisch, S., Rueden, C., Saalfeld, S., Schmid, B., Tinevez, J.-Y., White, D.J., Hartenstein, V., Eliceiri, K., Tomancak, P. and Cardona, A. (2012) Fiji: an open-source platform for biological-image analysis Nature Methods 9, 676–682.

26. Deshpande, V., Luebeck, J., Nguyen, N.D., Bakhtiari, M., Turner, K.M., Schwab, R., Carter, H., Mischel, P.S. and Bafna, V. (2019) Exploring the landscape of focal amplifications in cancer using AmpliconArchitect. Nat Commun 10, 392.

27. Ryba, T., Battaglia, D., Pope, B.D., Hiratani, I. and Gilbert, D.M. (2011) Genome-scale analysis of replication timing: from bench to bioinformatics. Nat Protoc 6, 870–895.

28. Li, H., Handsaker, B., Wysoker, A., Fennell, T., Ruan, J., Homer, N., Marth, G., Abecasis, G., Durbin, R. and 1000, G.P.D.P.S. (2009) The Sequence Alignment/Map format and SAMtools. Bioinformatics 25, 2078–2079.

29. Robinson, J.T., Thorvaldsdóttir, H., Winckler, W., Guttman, M., Lander, E.S., Getz, G. and Mesirov, J.P. (2011) Integrative genomics viewer. Nat Biotechnol 29, 24–26.

30. Botta-Dukát, Z. (2005) Rao’s quadratic entropy as a measure of functional diversity based on multiple traits Journal of Vegetation Science 16, 533–540.

31. Rao, C.R. (1980) Diversity and dissimilarity coefficients: A unified approach Theoretical Population Biology 21, 24–43.

32. Shannon, C.E. (1948) A mathematical theory of communication The Bell System Technical Journal 27, 379–423.

33. Jones, M.J.K., Rai, S.K., Pfuderer, P.L., Bonfim-Melo, A., Pagan, J.K., Clarke, P.R., McClelland, S.E. and Boemo, M.A. (2022) A high-resolution, nanopore-based artificial intelligence assay for DNA replication stress in human cancer cells bioRxiv 4.

34. Quinn, L.A., Moore, G.E., Morgan, R.T. and Woods, L.K. (1979) Cell lines from human colon carcinoma with unusual cell products, double minutes, and homogeneously staining regions Cancer Research 39, 4914–4924.

35. Gu, Z., Gu, L., Eils, R., Schlesner, M. and Brors, B. (2014) circlize Implements and enhances circular visualization in R. Bioinformatics 30, 2811–2812.

36. Bedrat, A., Lacroix, L. and Mergny, J.L. (2016) Re-evaluation of G-quadruplex propensity with G4Hunter. Nucleic Acids Res 44, 1746–1759.

37. Bodnar, A.G., Ouellette, M., Frolkis, M., Holt, S.E., Chiu, C.-P., Morin, G.B., Harley, C.B., Shay, J.W., Lichtsteiner, S. and Wright, W.E. (1998) Extension of life-span by introduction of telomerase into normal human cells Science 279, 349–352.

38. Luebeck, J., Coruh, C., Dehkordi, S.R., Lange, J.T., Turner, K.M., Deshpande, V., Pai, D.A., Zhang, C., Rajkumar, U., Law, J.A., Mischel, P.S. and Bafna, V. (2020) AmpliconReconstructor integrates NGS and optical mapping to resolve the complex structures of focal amplifications. Nat Commun 11, 4374.

39. Hung, K.L., Luebeck, J., Dehkordi, S.R., Colón, C.I., Li, R., Wong, I.T., Coruh, C., Dharanipragada, P., Lomeli, S.H., Weiser, N.E., Moriceau, G., Zhang, X., Bailey, C., Houlahan, K.E., Yang, W., González, R.C., Swanton, C., Curtis, C., Jamal-Hanjani, M., Henssen, A.G., Law, J.A., Greenleaf, W.J., Lo, R.S., Mischel, P.S., Bafna, V. and Chang, H.Y. (2022) Targeted profiling of human extrachromosomal DNA by CRISPR-CATCH. Nat Genet 54, 1746–1754.

40. Dileep, V. and Gilbert, D.M. (2018) Single-cell replication profiling to measure stochastic variation in mammalian replication timing. Nat Commun 9, 427.

41. Rivera-Mulia, J.C. and Gilbert, D.M. (2016) Replication timing and transcriptional control: beyond cause and effect-part III. Curr Opin Cell Biol 40, 168–178.

42. Vassilev, L. and Johnson, E.M. (1990) An initiation zone of chromosomal DNA replication located upstream of the c-myc gene in proliferating HeLa cells. Mol Cell Biol 10, 4899–4904.

43. Akerman, I., Kasaai, B., Bazarova, A., Sang, P.B., Peiffer, I., Artufel, M., Derelle, R., Smith, G., Rodriguez-Martinez, M., Romano, M., Kinet, S., Tino, P., Theillet, C., Taylor, N., Ballester, B. and Méchali, M. (2020) A predictable conserved DNA base composition signature defines human core DNA replication origins. Nat Commun 11, 4826.

44. Guilbaud, G., Murat, P., Wilkes, H.S., Lerner, L.K., Sale, J.E. and Krude, T. (2022) Determination of human DNA replication origin position and efficiency reveals principles of initiation zone organisation. Nucleic Acids Res 50, 7436–7450.

45. Ge, X.Q., Jackson, D.A. and Blow, J.J. (2007) Dormant origins licensed by excess Mcm2-7 are required for human cells to survive replicative stress. Genes Dev 21, 3331–3341.

46. Ibarra, A., Schwob, E. and Méndez, J. (2008) Excess MCM proteins protect human cells from replicative stress by licensing backup origins of replication Proceedings of the National Academy of Sciences 105, 8956–8961.

47. Hyrien, O., Guilbaud, G. and Krude, T. (2025) The double life of mammalian DNA replication origins. Genes Dev 39, 304–324.

48. Pope, B.D., Ryba, T., Dileep, V., Yue, F., Wu, W., Denas, O., Vera, D.L., Wang, Y., Hansen, R.S., Canfield, T.K., Thurman, R.E., Cheng, Y., Gülsoy, G., Dennis, J.H., Snyder, M.P., Stamatoyannopoulos, J.A., Taylor, J., Hardison, R.C., Kahveci, T., Ren, B. and Gilbert, D.M. (2014) Topologically associating domains are stable units of replication-timing regulation. Nature 515, 402–405.

49. Petermann, E., Orta, M.L., Issaeva, N., Schultz, N. and Helleday, T. (2010) Hydroxyurea-stalled replication forks become progressively inactivated and require two different RAD51-mediated pathways for restart and repair. Mol Cell 37, 492–502.

50. Ilić, M., Zaalberg, I.C., Raaijmakers, J.A. and Medema, R.H. (2022) Life of double minutes: generation, maintenance, and elimination. Chromosoma 131, 107–125.

51. Helmrich, A., Ballarino, M. and Tora, L. (2011) Collisions between replication and transcription complexes cause common fragile site instability at the longest human genes. Mol Cell 44, 966–977.

