## Supplementary Figures for "ecDNA replication is disorganised and vulnerable to replication stress"

**Supplementary Figure S1**

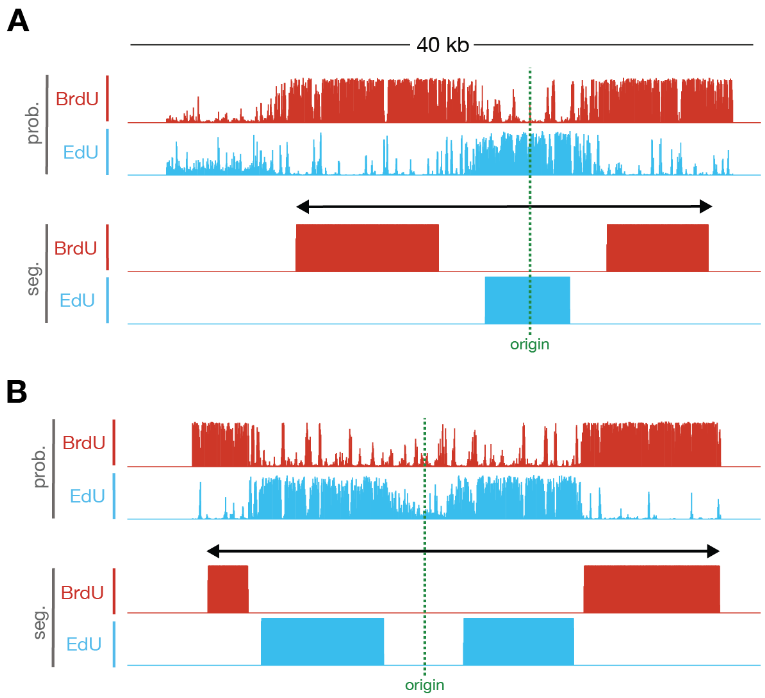


**Supplementary Figure S1: Origins on ecDNA from FACS isolation protocol.** Representative raw base analogue incorporation probabilities and DNAscent segmentation of origins of replication each represented by four tracks (Upper tracks: Raw BrdU and EdU probabilities (prob.) at thymidine positions; Lower tracks BrdU and EdU segmentation (seg.) derived from the raw probabilities). Both origins map to ecDNA in the untreated COLO 320DM cell line (coordinates on ecDNA map chromosome chr8_126425747-127997820 for origin track on top are 41,538 to 81,538, origin track on bottom 1,251,000 to 1,291,000.

### Supplementary Figure S2

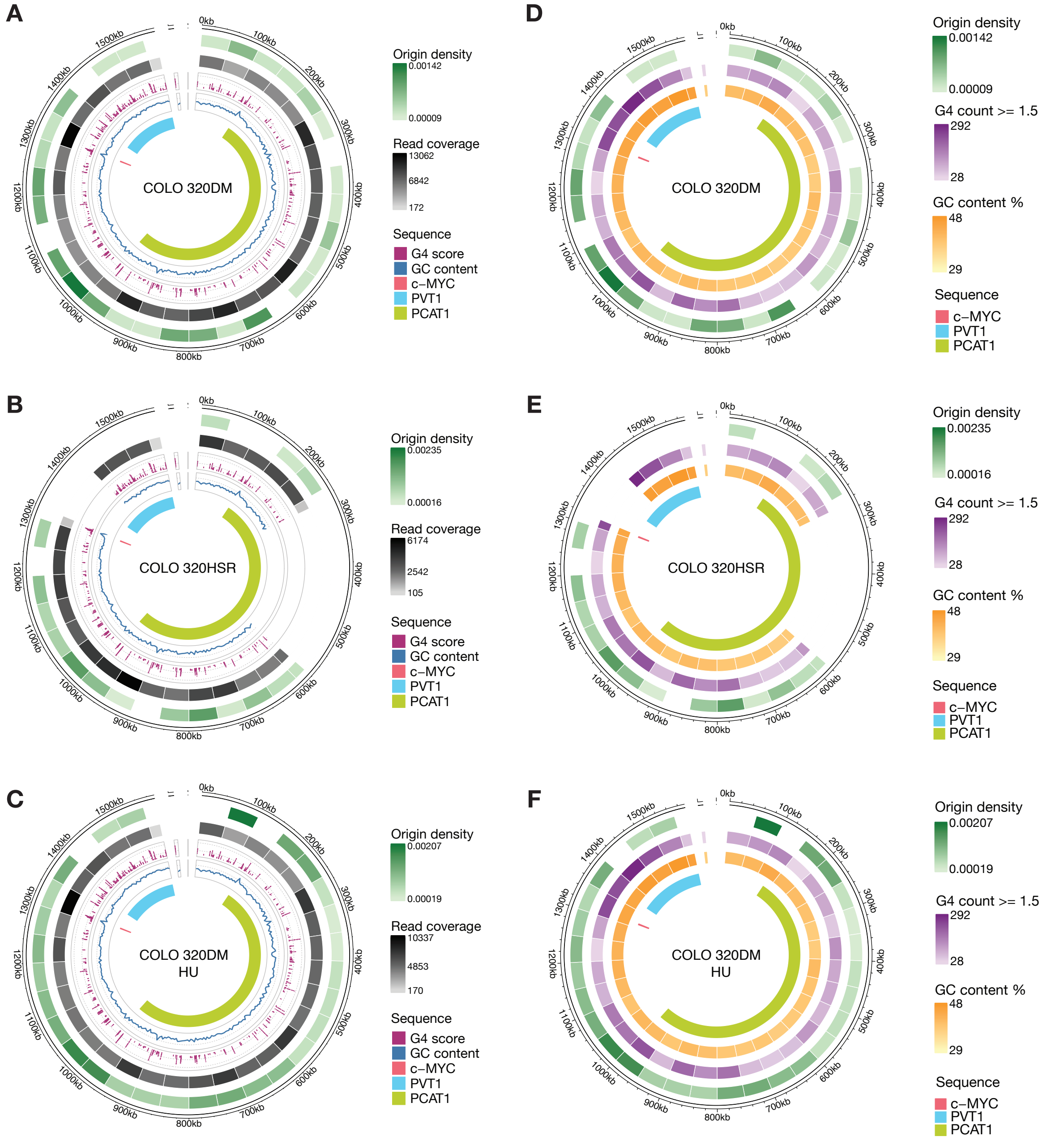


**Supplementary Figure S2: Origin densities vs. read coverage, GC density and G4 count. A. – C.** Origin density and read coverage averaged over 50 kb windows of the Colo 320DM ecDNA interval in **A.** Colo 320DM, **B.** Colo 320HSR and **C.** Colo 320DM treated with HU. G4 calls from G4Hunter (1) above threshold 1.5 (purple dots); grey dotted lines are shown at G4Hunter scores 2.0 and 3.0); GC content (blue line); grey dotted line shown at 50%) and key genes: c-*MYC* (salmon), *PVT1* (light blue) and *PCAT1* light green). **D. – F.** Origin densities show alongside averaged G4 count (G4s with a G4Hunter (1) score >= 1.5), and GC content in 50 kb segments in **D.** Colo 320DM, **E.** Colo 320HSR and **F.** Colo 320DM treated with HU.

**Supplementary Figure S3**

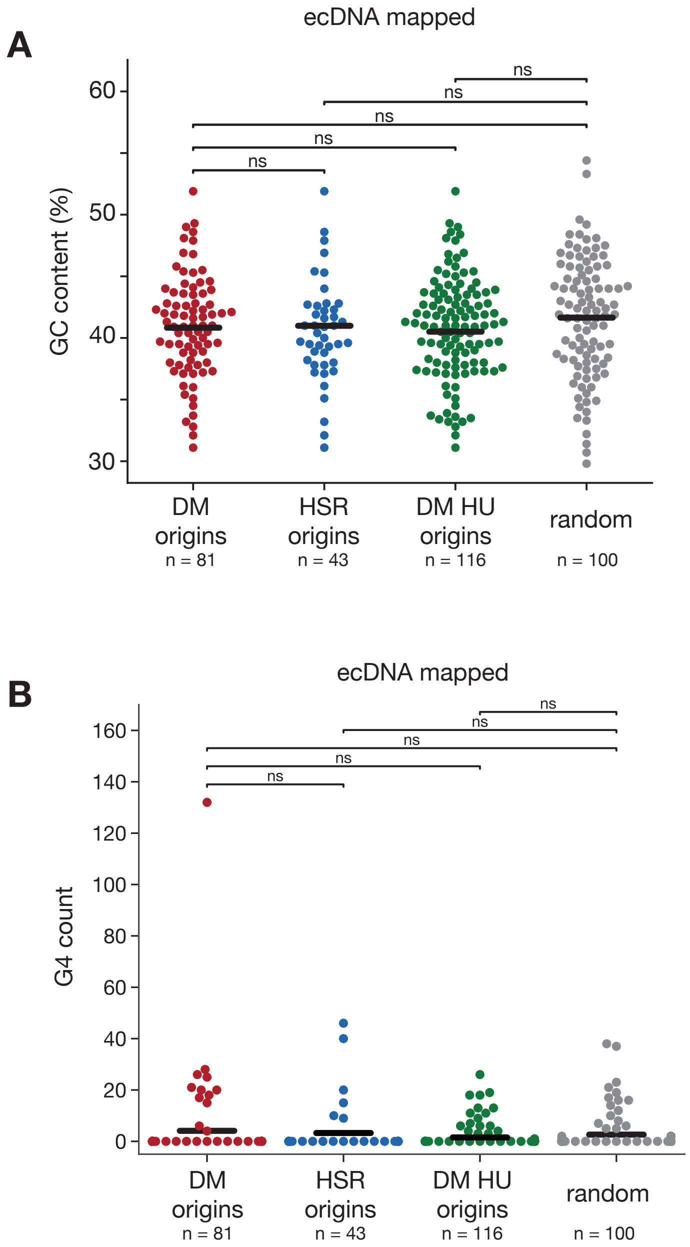


**Supplementary Figure S3. Quantitation of GC content and G4 count at DNAscent-detected origins in the COLO 320DM ecDNA interval.** GC content of all origin regions identified in the ecDNA interval compared to 100 randomly selected non-origin regions in the ecDNA interval. **A.** GC content of all origins on ecDNA, with origin size set to 1kb around the midpoint of the origin coordinate in COLO 320DM (red, mean = 40.67), COLO 320HSR (blue, mean = 40.51) and HU-treated COLO 320DM (green, mean = 40.73) compared to 100 randomly selected non-origin regions (grey, mean = 41.51) of the same size on ecDNA. **B.** G4 count (G4s with G4Hunter score >= 1.5) at of all origins identified in COLO 320DM (red, mean = 4.099), COLO 320HSR (blue, mean = 3.256) and HU-treated COLO 320DM (green, mean = 1.534) in the ecDNA interval compared to 100 randomly selected non-origin regions (grey, mean = 2.630) in the ecDNA interval. There is one outlier in the DM origins (coordinates 980,300 to 981,300 on chr8_126425747-127997820 of the ecDNA reference map), where G4 Hunter calls 132 G4s. Please note, that this is the maximum number of possible G4-forming sequence patterns detected and as they are all subsequently called, it is unlikely that 132 G4s would form within this 1,000 bp origin segment, P-values are obtained from a two-sided non-parametric Wilcoxon Rank Sum test. Statistical significance: ns = not significant ($p \geq 0.05$).

**Supplementary Figure S4**

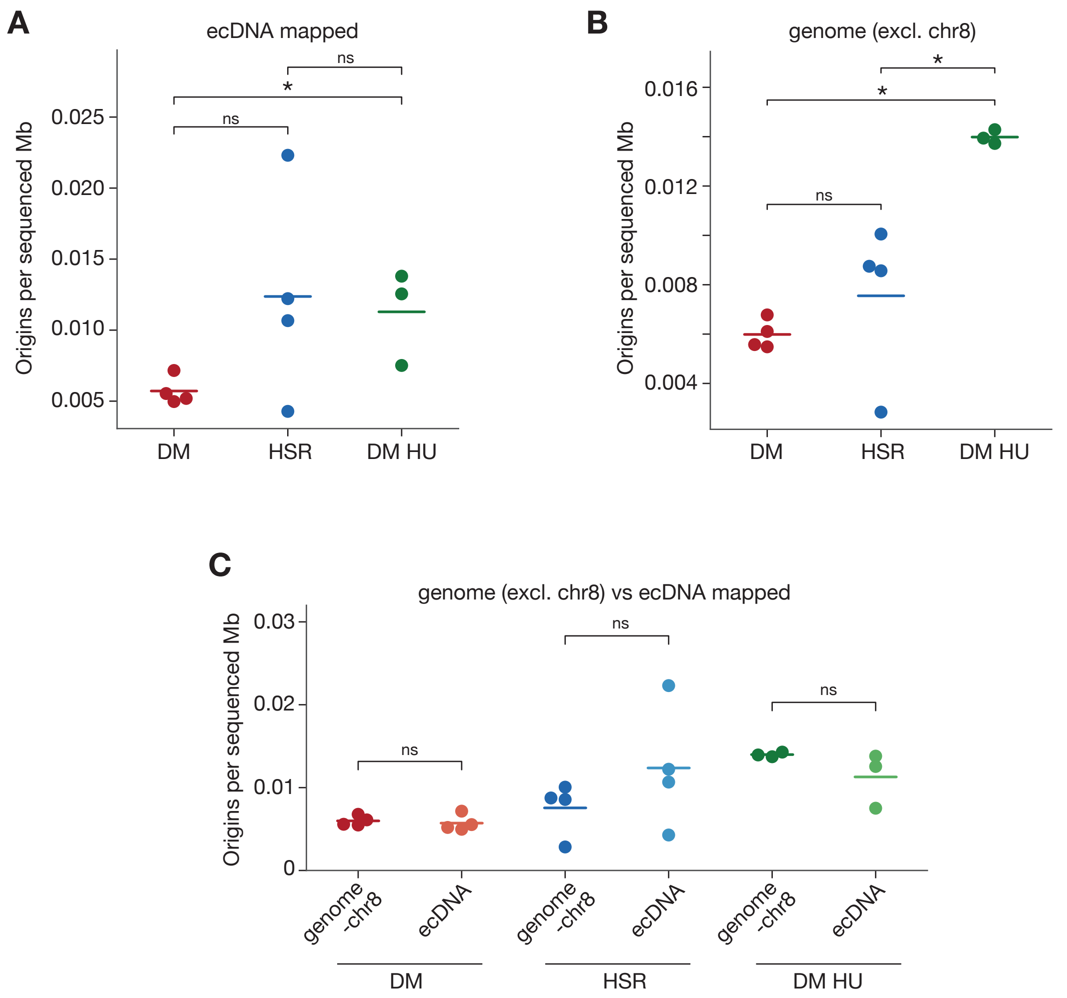


**Supplementary Figure S4: Origin densities on ecDNA. A.** Origin firing rates on ecDNA in Colo 320DM (red, mean = 0.0057), Colo 320HSR (blue, mean = 0.0124) and Colo 320DM treated with HU (green, mean = 0.0113) as indicated by origins per sequenced mega base (mb). **B.** Origin firing rates on whole genome mapped reads (excluding chromosome 8) in Colo 320DM (red, mean = 0.006), Colo 320HSR (blue, mean = 0.0073) and Colo 320DM treated with HU (green, mean = 0.0141) as indicated by origins per sequenced mega base (mb) **C.** Side-by-side comparison of origins in the genome (except chr 8) and the ecDNA interval in each line / condition. The same data is plotted as in A & B. All p-values are obtained from a two-sided non-parametric Wilcoxon Rank Sum test. Statistical significance: ns = not significant ($p \geq 0.05$), * $p < 0.05$.

### Supplementary Figure S5

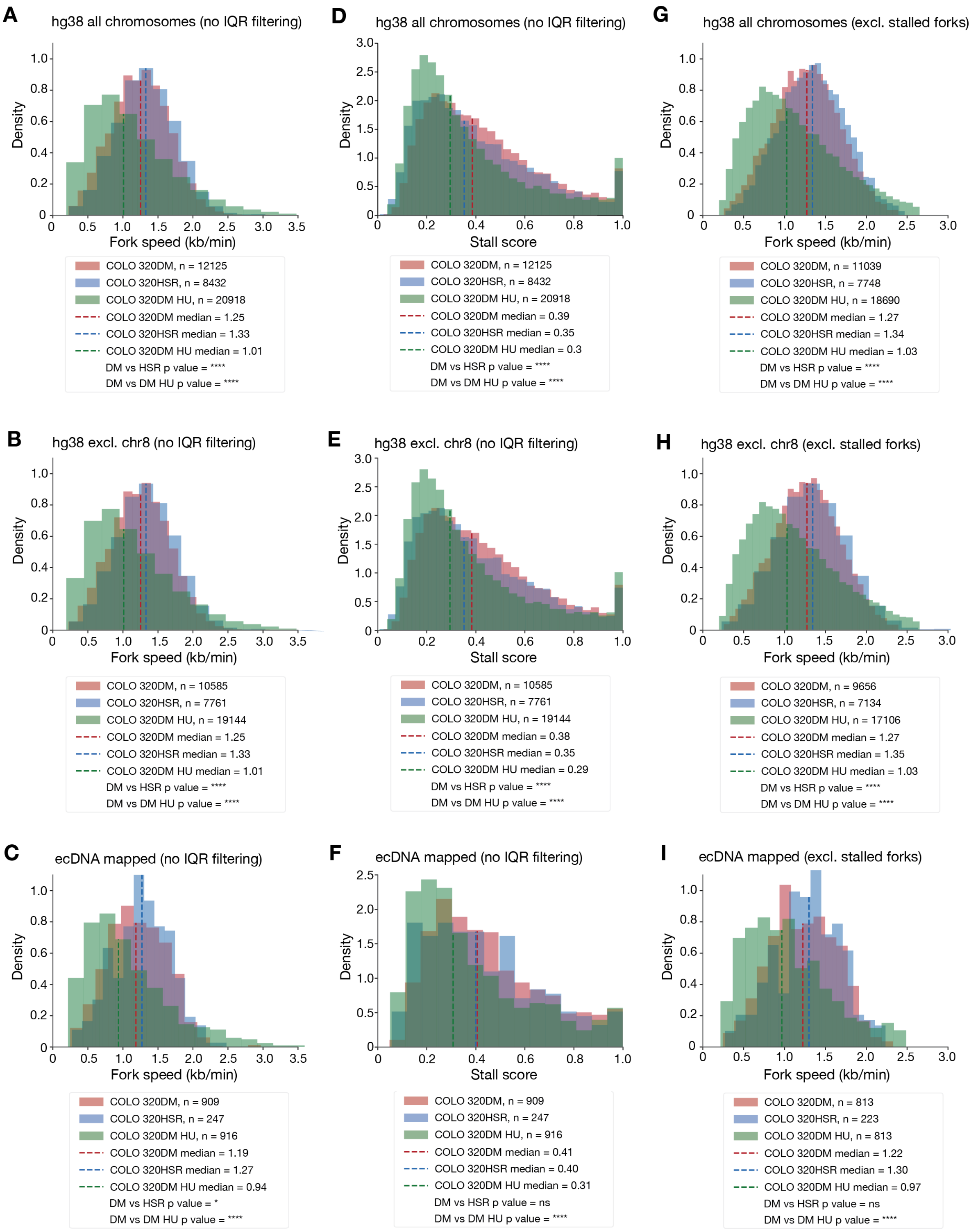


**Supplementary Figure S5. Raw fork speeds and stall scores for all chromosomes, or ecDNA reference.** Fork speed without interquartile range (IQR) filtering forks mapped to **A.** all chromosomes (hg38), **B.** all chromosomes excluding chromosome 8 and **C.** the ecDNA reference map. Stall scores without IQR filtering for forks mapped to **D.** all chromosomes, **E.** all chromosomes excluding chromosome 8 and **F.** the ecDNA reference map. Fork speeds excluding forks with a stall score of >= 0.8 mapped to **G.** all chromosomes, **H.** all chromosomes excluding chromosome 8 and **I.** the ecDNA reference map. All p-values for fork speeds are obtained from a two-sided Welch's t-test with no assumption of equal variances and all p-values for stall scores are obtained from a two-sided non-parametric Wilcoxon Rank Sum test. Statistical significance: ns = not significant ($p \geq 0.05$), * $p < 0.05$, ** $p < 0.01$, *** $p < 0.001$, **** $p < 0.0001$.

**Supplementary Figure S6**

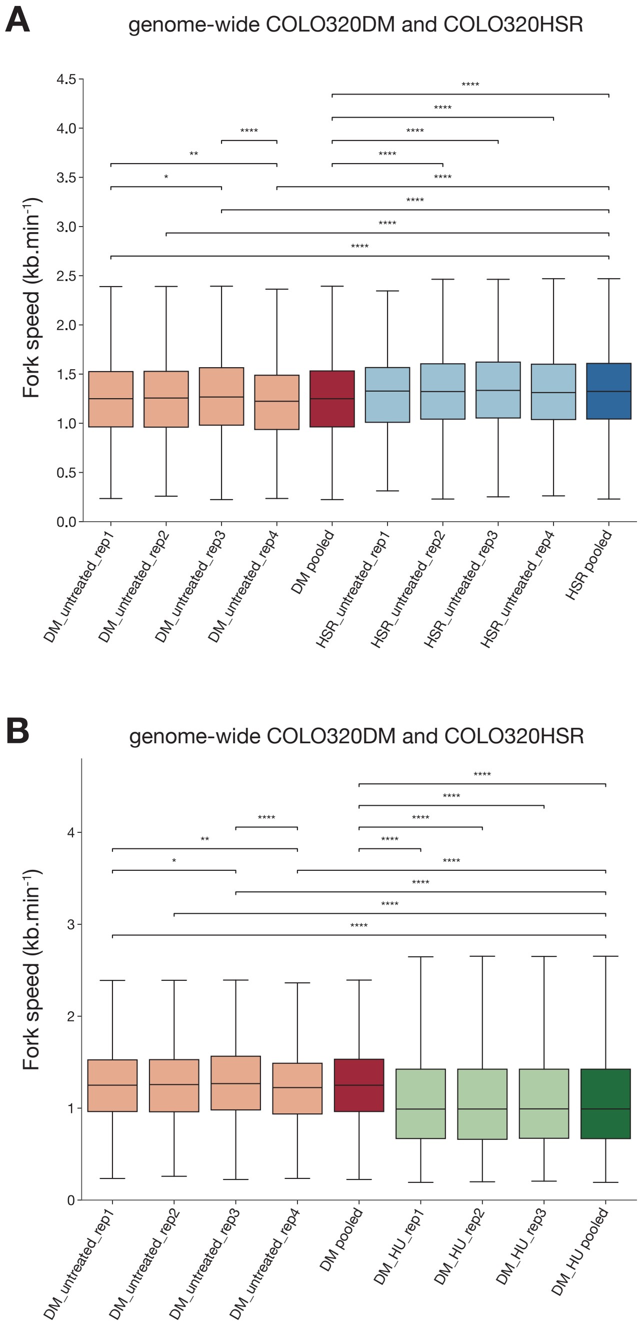


**Supplementary Figure S6. Fork speed comparison of individual replicates from PromethION runs. A.** Comparison of COLO 320DM samples untreated (red) vs COLO 320HSR (blue) replicates. B. Comparison of COLO 320DM samples untreated vs HU-treated COLO 320DM replicates (green). All p-values are obtained from a two-sided Welch's t-test with no assumption of equal variances. Statistical significance: ns = not significant ($p \geq 0.05$), * $p < 0.05$, ** $p < 0.01$, *** $p < 0.001$, **** $p < 0.0001$.

**Supplementary Figure S7**

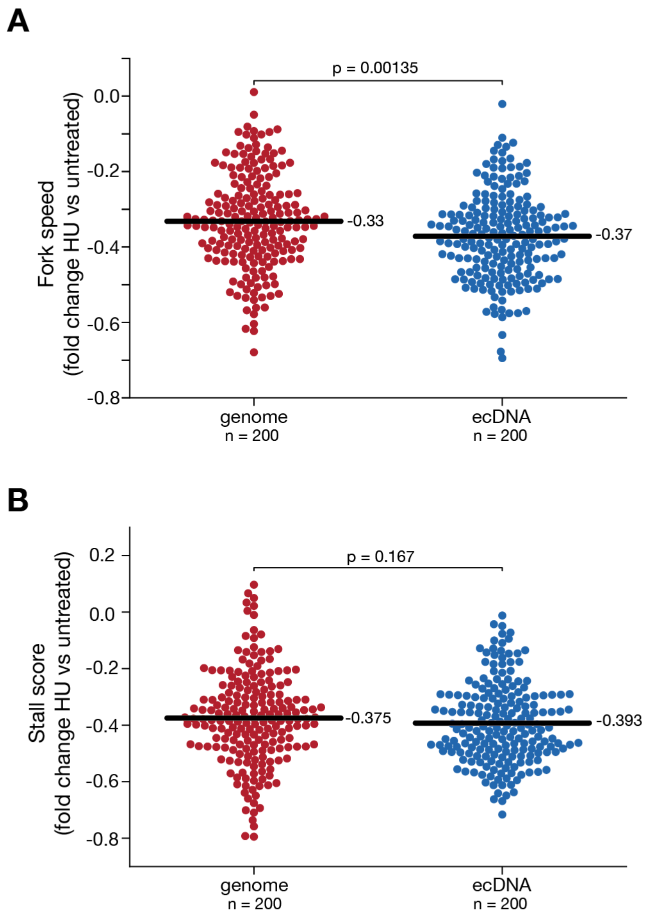


**Supplementary Figure S7. Differential effect of HU treatment on ecDNA compared to chromosomal DNA. A.** Effect of HU treatment on replication fork speed measured as log fold change (HU treated / untreated). Each datapoint represents the log fold change for a comparison between fork speed medians for 100 randomly sampled replication forks from the genome (chromosomes excluding chromosome 8, red) to ecDNA (blue) dataset, repeated 200 times. **B.** Same procedure as in A. but for replication fork stall scores. The mean for each experiment is shown as a black horizontal bar. P-values are obtained from a two-sided non-parametric Wilcoxon Rank Sum test.

**Supplementary Table S1.** Overview of biological replicates and datasets for method development. Enrichment strategy is only noted for samples included in Figure 1F, as well as the respective enrichment ratio. N50, mean and median read length are reported for reads that meet the minimum quality requirements we set when running DNAscent (minimum mapping length of 20kb, minimum mapping quality of 20). Number of detected replication forks is reported before and after filtering, with filters for replication forks requiring a minimum of 2kb spacing to the end of the mapped read, a stall score being calculated and passing the IQR-filter. Replication forks that are part of origin of replication or termination events are excluded from all fork counts.

### Supplementary References


1. Bedrat,A., Lacroix,L. and Mergny,J.L. (2016) Re-evaluation of G-quadruplex propensity with G4Hunter. *Nucleic Acids Res* **44**, 1746-1759.
